# The funny current I_f_ is essential for the fight-or-flight response in cardiac pacemaker cells

**DOI:** 10.1101/2022.05.07.490938

**Authors:** Colin H Peters, Christian Rickert, Stefano Morotti, Eleonora Grandi, Catherine Proenza

## Abstract

The sympathetic nervous system fight-or-flight response is characterized by a rapid increase in heart rate, which is mediated by an increase in the spontaneous action potential (AP) firing rate of pacemaker cells in the sinoatrial node. Sympathetic neurons stimulate sinoatrial myocytes (SAMs) by activating β adrenergic receptors (βARs) and increasing cAMP. The funny current (I_f_) is among the cAMP-sensitive currents in SAMs. I_f_ is critical for pacemaker activity, however, its role in the fight-or-flight response remains controversial. In this study, we used AP waveform analysis, machine learning, and dynamic clamp experiments in acutely-isolated SAMs from mice to quantitatively define the AP waveform changes and role of I_f_ in the fight-or-flight response. We found that while βAR stimulation significantly altered nearly all AP waveform parameters, the increase in AP firing rate was only correlated with changes in a subset of parameters (diastolic duration, late AP duration, and diastolic depolarization rate). Dynamic clamp injection of the βAR-sensitive component of I_f_ showed that it accounts for approximately 41% of the fight-or-flight increase in AP firing rate and 60% of the decrease in the interval between APs. Thus, I_f_ is an essential contributor to the fight-or-flight increase in heart rate.

## Introduction

The sympathetic fight-or-flight response is characterized by a rapid increase in heart rate that is mediated by pacemaker cells in the sinoatrial node. Sinoatrial node myocytes (SAMs) trigger each beat of the heart by firing spontaneous action potentials (APs) that propagate through the cardiac conduction system to initiate contraction (Keith and Flack, 1907). The sympathetic nervous system increases heart rate by accelerating this spontaneous AP firing rate. Within SAMs, the faster AP firing rate is caused by an increase in cytosolic cAMP following sympathetic activation of β adrenergic receptors (βARs) on the cell membrane. While a host of processes in SAMs are regulated directly and indirectly by the βAR-mediated increase in cAMP, the relative contributions of these processes to the fight-or-flight response remain unresolved.

APs in sinoatrial myocytes have a characteristic shape, which is determined by the unique complement of ion channels and transporters on the cell membrane. The changes to the sinoatrial AP waveform that occur with βAR stimulation could, in theory, serve as “fingerprints” to identify the ionic currents that mediate the increase in AP firing rate. However, the changes to the sinoatrial AP that occur with βAR stimulation have yet to be systematically defined, owing in part to 1) differences in experimental conditions and a lack of standard AP waveform definitions, which restrict comparisons between studies; 2) a focus on the diastolic depolarization rate (DDR) as the primary determinant of firing rate to the exclusion of other AP waveform parameters; and 3) the limited resolution of most studies owing to the heterogeneity of sinoatrial APs and difficulty of recording from large numbers of isolated SAMs.

In particular, whether the hyperpolarization-activated funny current, I_f_, contributes to the fight-or-flight response is the subject of longstanding debate (Lakatta and DiFrancesco, 2009; Bucchi et al., 2012; Hennis et al., 2021). I_f_ is a hallmark of SAMs and is critical for the generation of spontaneous APs (Bucchi et al., 2007; DiFrancesco, 2010; Baruscotti et al., 2011). I_f_ is produced by hyperpolarization-activated cyclic nucleotide-sensitive (HCN) channels, particularly HCN4, which is the predominant isoform in the sinoatrial node (Moosmang et al., 1999). Reduction in I_f_ via pharmacological blockers or mutations in HCN4 slows AP firing rate in SAMs and heart rate in human patients and animal models (Bucchi et al., 2007; Swedberg et al., 2010; Baruscotti et al., 2011; Milano et al., 2014). And cAMP potentiates I_f_, consistent with a role in the fight-or-flight response (Brown et al., 1979). However, multiple groups have argued that I_f_ plays only a minor role owing in part to its low fractional activation at physiological potentials in SAMs and a residual βAR response in HCN4 knockout animals (Lakatta and DiFrancesco, 2009; Alig et al., 2009; Mesirca et al., 2014; Hennis et al., 2021).

We recently showed that I_f_ is persistently active throughout the entire AP in SAMs (Peters et al., 2021). Consequently, I_f_ conducts more than 50% of the net inward charge movement during diastole, despite its low fractional activation (Peters et al., 2021). Importantly, the fraction of the net inward charge movement conducted by I_f_ increases to 93% of the total during βAR stimulation (Peters et al., 2021). This result strongly suggests that I_f_ contributes substantially to the fight-or-flight increase in heart rate.

In this study, we sought to quantitatively evaluate the role of I_f_ in the fight-or-flight response in SAMs. We began by analyzing a large data set of sinoatrial APs with automated AP waveform analysis, correlation analysis, and machine learning to identify the subset of AP waveform parameters that account for the majority of the βAR-mediated increase in AP firing rate. We then used dynamic clamp experiments to demonstrate that βAR-stimulation of I_f_ alone accounts for approximately 41%of the chronotropic response to βAR stimulation in SAMs, mainly via effects during diastole. These results clearly establish that I_f_ contributes to the fight-or-flight response as part of a multifaceted cardiac pacemaking mechanism.

## Results

### Most sinoatrial AP waveform parameters change in response to βAR stimulation

To define the effects of βAR stimulation on the sinoatrial AP waveform, we recorded spontaneous APs in the amphotericin perforated-patch configuration in 50 sinoatrial myocytes from 24 mice. In each cell, APs were first recorded in a control extracellular solution containing 1 nM of the βAR agonist isoproterenol (Iso) and then upon wash-on of 1 µM Iso. 1 nM Iso was used in the control solution as in previous studies (Larson et al., 2013; Peters et al., 2021) because it mimics the resting catecholamine levels in mice (Pichavaram et al., 2018; Lucot et al., 2005) and humans (Messan et al., 2017). We confirmed that 1 nM Iso did not increase AP firing rate compared to a solution lacking Iso (304 ± 15 bpm in 1 nM, N = 50 vs. 313 ± 31 bpm in 0 nM, N = 19; P = 0.9879). 1 µM Iso was used as the stimulated condition as previously (Larson et al., 2013; Fenske et al., 2020; Peters et al., 2021) and we confirmed 1 µM Iso to be a saturating concentration, as 10 µM Iso did not produce a further increase in AP firing rate (400 ± 12 bpm in 1 µM, N = 50 vs. 400 ± 33 bpm in 10 µM, N = 9; P = 1.0000).

AP firing rate and AP waveform parameters in the control and Iso-stimulated conditions were determined by automated analysis with ParamAP software (Rickert and Proenza, 2017, 2021). ParamAP defines 15 AP waveform parameters, including time intervals, membrane potentials, and rates of change of membrane potential, which together provide a full description of sinoatrial APs (**Table 1, Fig. S1**). Although AP waveforms recorded in individual cells exhibited considerable heterogeneity (**Fig. 1A** and **B**), characteristic features of sinoatrial APs were apparent in the recordings, including a spontaneous diastolic depolarization, depolarized maximum diastolic potential (MDP), slow maximum upstroke velocity (MUV), and small AP amplitude (APA) (**Table 2**).

**Table 1:**
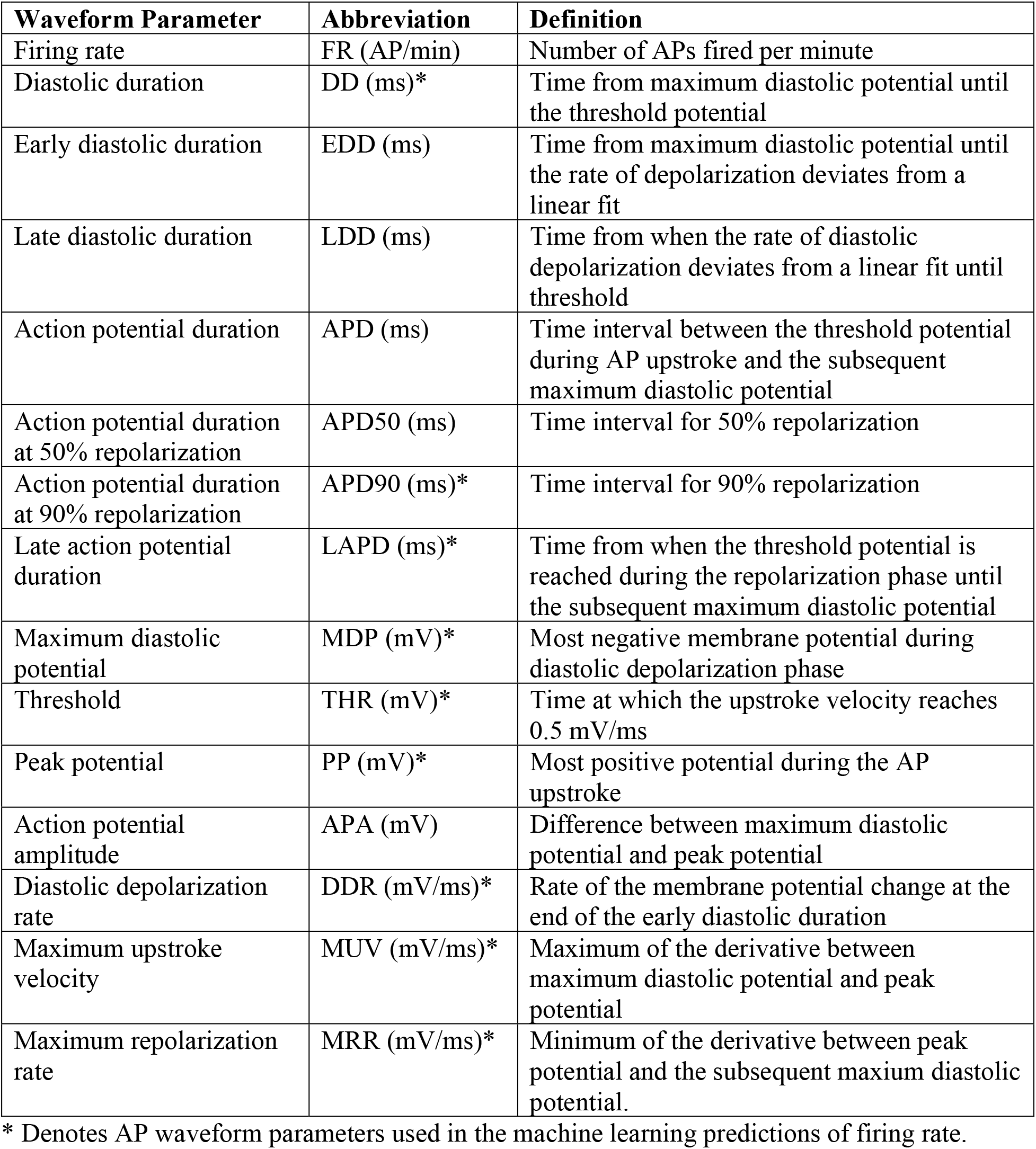
AP Waveform Parameter Definitions.

| Waveform Parameter | Abbreviation | Definition |
| --- | --- | --- |
| Firing rate | FR (AP/min) | Number of APs fired per minute |
| Diastolic duration | DD (ms)* | Time from maximum diastolic potential until the threshold potential |
| Early diastolic duration | EDD (ms) | Time from maximum diastolic potential until the rate of depolarization deviates from a linear fit |
| Late diastolic duration | LDD (ms) | Time from when the rate of diastolic depolarization deviates from a linear fit until threshold |
| Action potential duration | APD (ms) | Time interval between the threshold potential during AP upstroke and the subsequent maximum diastolic potential |
| Action potential duration at 50% repolarization | APD50 (ms) | Time interval for 50% repolarization |
| Action potential duration at 90% repolarization | APD90 (ms)* | Time interval for 90% repolarization |
| Late action potential duration | LAPD (ms)* | Time from when the threshold potential is reached during the repolarization phase until the subsequent maximum diastolic potential |
| Maximum diastolic potential | MDP (mV)* | Most negative membrane potential during diastolic depolarization phase |
| Threshold | THR (mV)* | Time at which the upstroke velocity reaches 0.5 mV/ms |
| Peak potential | PP (mV)* | Most positive potential during the AP upstroke |
| Action potential amplitude | APA (mV) | Difference between maximum diastolic potential and peak potential |
| Diastolic depolarization rate | DDR (mV/ms)* | Rate of the membrane potential change at the end of the early diastolic duration |
| Maximum upstroke velocity | MUV (mV/ms)* | Maximum of the derivative between maximum diastolic potential and peak potential |
| Maximum repolarization rate | MRR (mV/ms)* | Minimum of the derivative between peak potential and the subsequent maximum diastolic potential. |

\* Denotes AP waveform parameters used in the machine learning predictions of firing rate.

**Figure 1:**
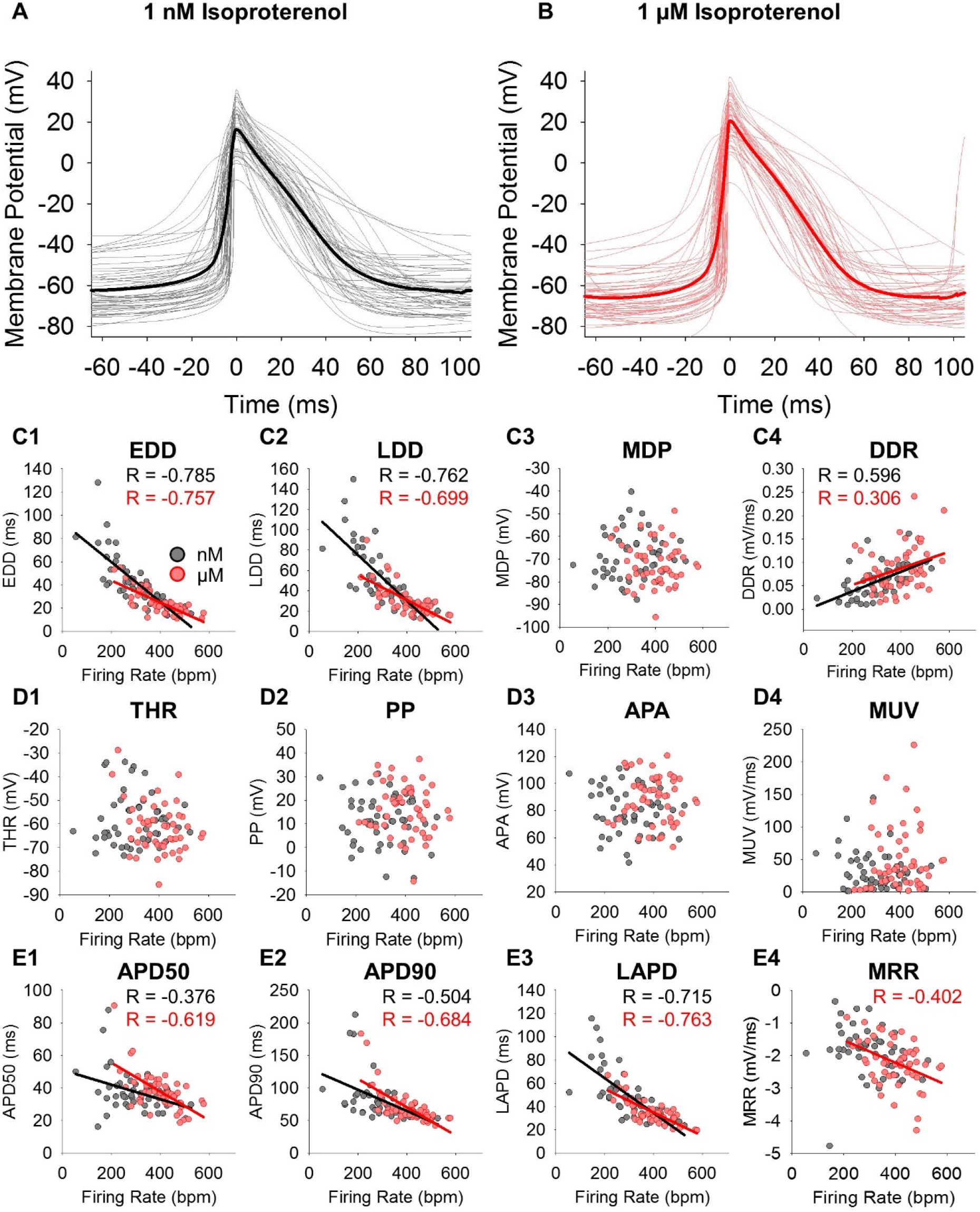
Sinoatrial Action Potentials are Heterogeneous. **A:** Overlays of individual (N=50) sinoatrial AP waveforms (*grey*) and the average of all waveforms (*black*) recorded in 1 nM Iso. **B:** Overlays of individual (N=50) sinoatrial AP waveforms (*pink*) and the average of all waveforms (*red*) recorded in 1 µM Iso. **C-E:** Correlation plots of individual AP waveform parameters in 1 nM Iso (*grey*) and 1 µM Iso (*pink*) as a function of the AP firing rate. Significant correlations are shown as black (1 nM) and red (1 µM) lines and the Pearson correlation coefficients are noted. All correlation coefficients and details of statistical tests can be found in **Table 2**.

**Table 2:** Average (± SEM) AP Firing Rate and Waveform Parameters in 1 nM and 1 µM Isoproterenol.

| | 1 nM Isoproterenol | 1 μM Isoproterenol | P-Value 1 nM vs. 1 μM |
| --- | --- | --- | --- |
| FR (bpm) | 304 ± 14 | 400 ± 12 | **P < 0.0001** |
| DD (ms) | 93.7 ± 7.1 | 55.9 ± 3.5 | **P < 0.0001** |
| EDD (ms) | 41.9 ± 3.1 | 25.0 ± 1.5 | **P < 0.0001** |
| LDD (ms) | 51.7 ± 4.3 | 30.9 ± 2.1 | **P < 0.0001** |
| MDP (mV) | -68.7 ± 1.4 | -72.0 ± 1.3 | **P = 0.0013** |
| DDR (mV/ms) | 0.060 ± 0.005 | 0.087 ± 0.007 | **P < 0.0001** |
| THR (mV) | -58.3 ± 1.5 | -61.9 ± 1.5 | **P = 0.0007** |
| PP (mV) | 11.9 ± 1.5 | 16.2 ± 1.5 | **P < 0.0001** |
| APA (mV) | 81.0 ± 2.5 | 88.3 ± 2.4 | **P < 0.0001** |
| MUV (mV/ms) | 32.6 ± 4.1 | 51.0 ± 7.0 | **P = 0.0007** |
| APD (ms) | 116.8 ± 5.1 | 99.6 ± 3.3 | **P < 0.0001** |
| APD50 (ms) | 37.4 ± 1.7 | 37.9 ± 1.7 | **P = 0.5651** |
| APD90 (ms) | 80.3 ± 4.7 | 70.5 ± 3.7 | **P = 0.0003** |
| LAPD (ms) | 48.8 ± 3.0 | 34.9 ± 1.6 | **P < 0.0001** |
| MRR (mV/ms) | -1.9 ± 0.1 | -2.2 ± 0.1 | **P < 0.0001** |

N = 50 for all cases
Red indicates a significant ( $P < 0.05$ ) correlation.

We performed correlation analysis to identify the AP waveform parameters that are significantly associated with AP firing rate (**Fig. 1C-E; Table S1**). We found that firing rate was significantly correlated with the duration of both early and late phases of the diastolic depolarization (EDD and LDD), with all measures of AP duration (APD), and with the diastolic depolarization rate (DDR) in both 1 nM and 1 µM Iso, while the maximum repolarization rate (MRR) was correlated with firing rate only in 1 µM Iso. Interestingly, firing rate was not significantly correlated with MDP or any other parameter reporting membrane voltage or with the upstroke velocity.

We next examined the changes in the AP waveform that occur in response to βAR stimulation. As expected, the AP firing rate was significantly greater in 1 µM Iso than in 1 nM Iso (**Table 2**; **Fig. 2B**). This increase in firing rate was accompanied by significant changes in every AP waveform parameter except APD50 (**Fig. 2B-E**). βAR stimulation shortened time intervals in the AP including the EDD, LDD, APD90, and late AP duration (LAPD). It also changed membrane potentials by hyperpolarizing the MDP and threshold (THR), depolarizing the peak potential (PP) and increasing the APA. Finally, βAR stimulation sped rates of change of membrane potential, increasing the DDR, MUV, and MRR.

**Figure 2:**
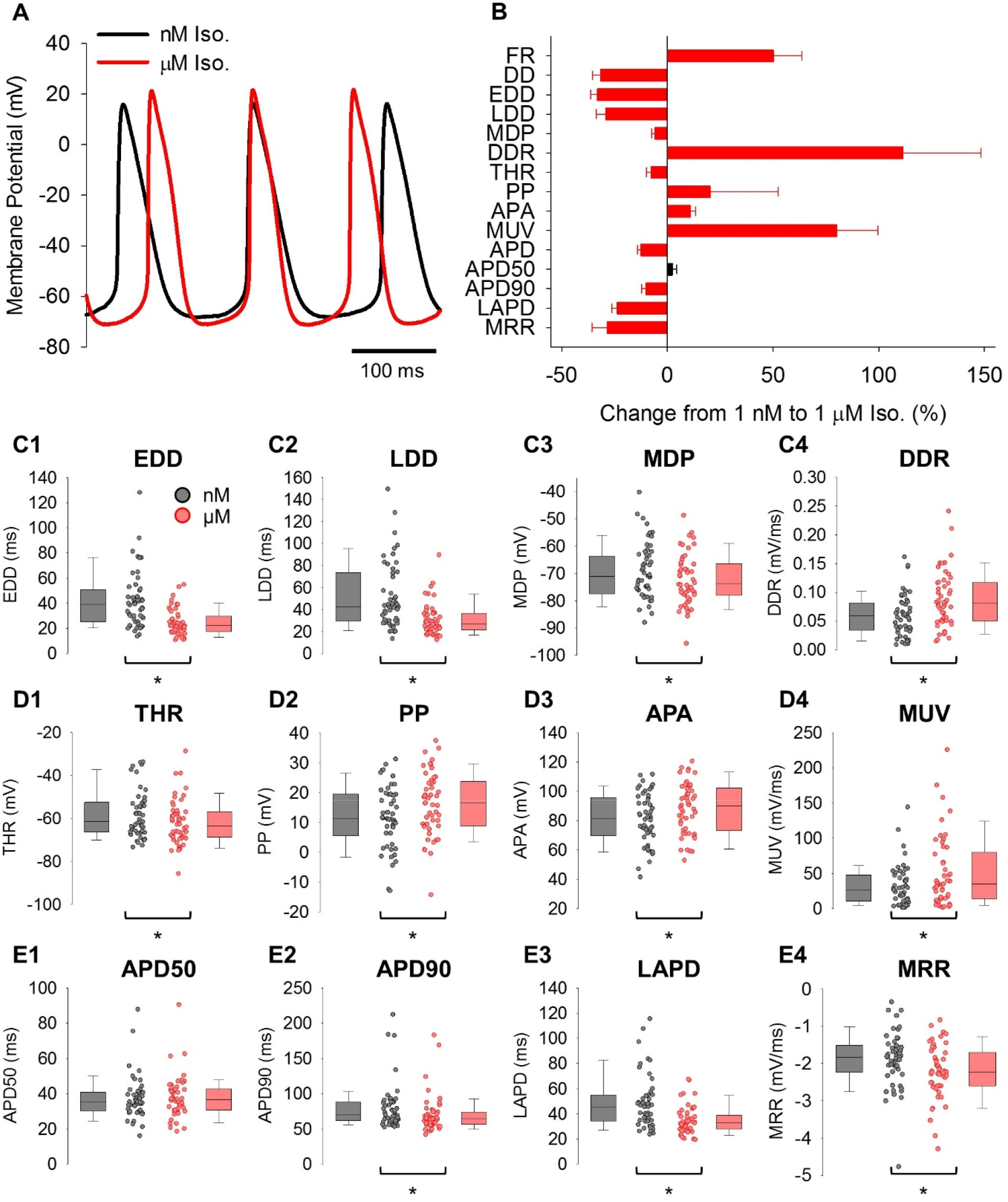
Isoproterenol Affects Most Sinoatrial AP Waveform Parameters. **A:** Representative sinoatrial APs recorded from a single cell in 1 nM Iso (*black*) and following wash-on of 1 µM Iso (*red*). **B:** Average (±SEM) percentage change in sinoatrial AP firing rate and waveform parameters induced by perfusion of 1 µM Iso. Red bars denote parameters that are significantly different after βAR stimulation. Since MDP and MRR are negative numbers, a negative percentage change is equivalent to hyperpolarization of the MDP and an increase in the magnitude of MRR. **C-E:** Box plots of median waveform parameters associated with the diastolic depolarization (**C1-C4**), action potential voltages and upstroke (**D1-D4)**, and action potential duration and repolarization (**E1-E4)** in 1 nM Iso (*grey*) and 1 µM Iso (*pink*). Individual recordings in **C-E** are shown as circles. Asterisks indicate significant differences between 1 nM and 1 µM Iso. N values and details of statistical tests are found in **Table 2**.

Overall, these data are consistent with a coordinated response of many Iso-sensitive currents during βAR stimulation. However, they do not provide information about the relative importance of different AP parameters in determining the increase in firing rate with βAR stimulation.

### Diastolic depolarization rate and late action potential duration are the primary predictors of the βAR increase in firing rate

We used two independent approaches to identify the AP waveform parameters that are the most important determinants of the Iso-dependent increase in AP firing rate. First, we calculated Pearson correlation coefficients to assess how Iso-induced changes in each AP waveform parameter relate to the increase in firing rate (**Table S1; Fig. 3A-C**). Unsurprisingly, the increase in firing rate in response to 1 µM Iso was significantly correlated with decreases in the time intervals that make up the total cycle length, including the diastolic duration (DD) and the total APD (**Table S1**). Within the diastolic duration, decreases in both EDD and LDD were significantly associated with the increase in firing rate (**Fig. 2A1** and **A2**). However, of the parameters comprising the total APD, only a decrease in LAPD was significantly correlated with increased firing rate (**Fig. 3C1-C3**). The DDR was the only non-interval parameter that was significantly correlated with the increase in firing rate (**Fig. 3A4**).

**Figure 3:**
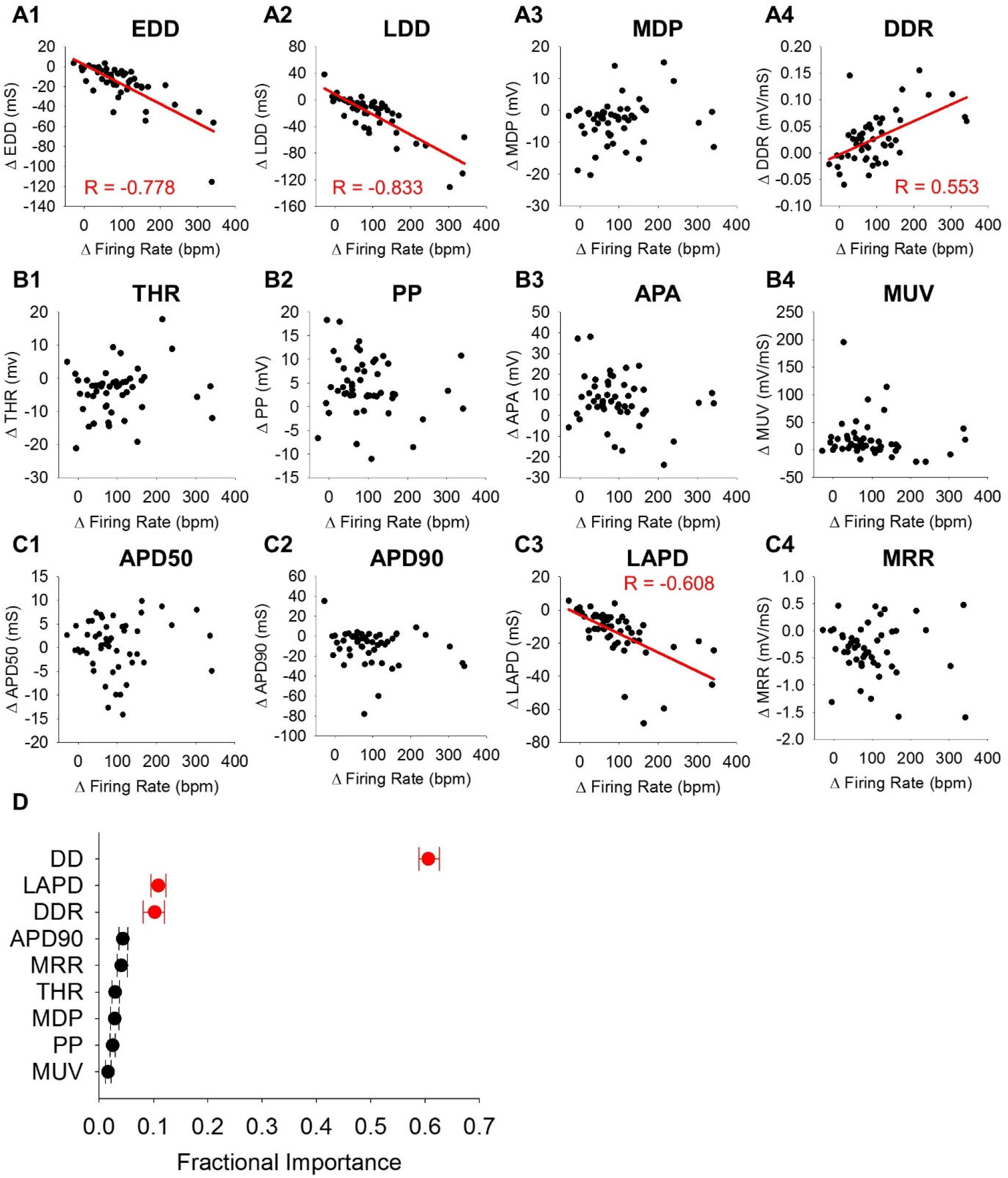
βAR Stimulation Accelerates AP Firing by Shortening the Diastolic Interval. Correlation plots of the change in waveform parameters associated with diastolic depolarization (**A1-A4**), action potential upstroke (**B1-B4)**, and action potential duration (**C1-E4)** versus the change in firing rate in SAMs between 1 nM and 1 µM Iso. Significant correlations are shown as a red line and the Pearson correlation coefficient is noted. All correlation coefficients and details of statistical tests can be found in **Supplementary Table S1. D:** Average importance of changes to waveform parameters during βAR stimulation in predicting concomitant changes in AP firing rate as measured by random forest machine learning analysis. Variables that were significantly correlated with changes in firing rate in **A**-**C** are shown in red. Error bars show the minimum and maximum values obtained from the 100 forests modelled.

We next used a machine learning approach as an independent means to identify the waveform parameters that are the most important predictors of the βAR-stimulated increase in firing rate. We used 100 random forests, each containing 1000 decision trees (Breiman, 2001), to evaluate the importance of a subset of waveform parameters. Consistent with the results of the correlation analysis, machine learning identified changes to DD, LAPD, and DDR as significantly associated with the βAR-induced increase in AP firing rate (**Fig. 3D**). Taken together, these results indicate that the interval between APs, including the LAPD, is the most important predictor of the increase in AP firing rate. It follows that the currents active at sub-threshold membrane potentials are the primary drivers of βAR-driven increases in sinoatrial node firing rate.

### Isoproterenol stimulation of I_f_ accounts for 41% of the βAR firing rate increase

Although many Iso-sensitive currents contribute to the diastolic depolarization, our previous study showed that Iso stimulation increases I_f_ from ∼50% to >90% of the net inward current during each AP (Peters et al., 2021). Therefore, we sought to determine the degree to which this increase in I_f_ accounts for the fight-or-flight increase in AP firing rate in SAMs, independent of the contribution of other Iso-sensitive currents. To this end, we employed the dynamic clamp approach using custom dynamic clamp hardware (Desai, 2017) and software (Rickert and Proenza, 2019a; b). Dynamic clamp is a hybrid experimental-computational method, in which mathematical models of currents are dynamically calculated based on instantaneous voltages measured from a patch-clamped cell and then injected back into the same cell in a fast (10 µs) feedback loop.

In these experiments, we defined the Iso-sensitive component of I_f_ (I_f-Iso_) in SAMs as the difference current between our previously published models of I_f_ in 1 µM and 1 nM Iso (Peters et al., 2021). In current-clamp experiments I_f-Iso_ was dynamically calculated from voltages measured in spontaneously-firing SAMs patch-clamped in the amphotericin perforated patch configuration. I_f-Iso_ was then either added to cells perfused with 1 nM Iso or subtracted from cells perfused with 1 µM Iso and the resulting changes in the AP firing rate and AP waveform parameters were compared those recorded in the same cell in the absence of a dynamic conductance. The overall approach is illustrated schematically in **Figure S2**.

As an initial proof of concept, spontaneously firing cells perfused with 1 nM Iso were dynamically injected with progressively increasing amounts of I_f-Iso_ with simulated maximal conductances ranging from 5-40 nS (**Fig. 4A**). We found that an average maximal I_f_ conductance of 10 nS was sufficient to significantly increase the AP firing rate and larger injections of I_f_ caused larger increases in firing rate (**Fig. 4C** and **4D**). In accordance, when the reverse experiment was performed and I_f-Iso_ was subtracted from SAMs perfused with 1 µM Iso, a 10 nS maximal I_f_ conductance was sufficient to significantly reduce firing rate (**Fig. 4B** and **4D**). Given our prior measurements showing an average maximum conductance of I_f_ of 11.79 ± 1.39 nS in SAMs (Peters et al., 2021), these results indicate that βAR stimulation of I_f_ contributes to the fight-or-flight increase in AP firing rate in SAMs.

**Figure 4:**
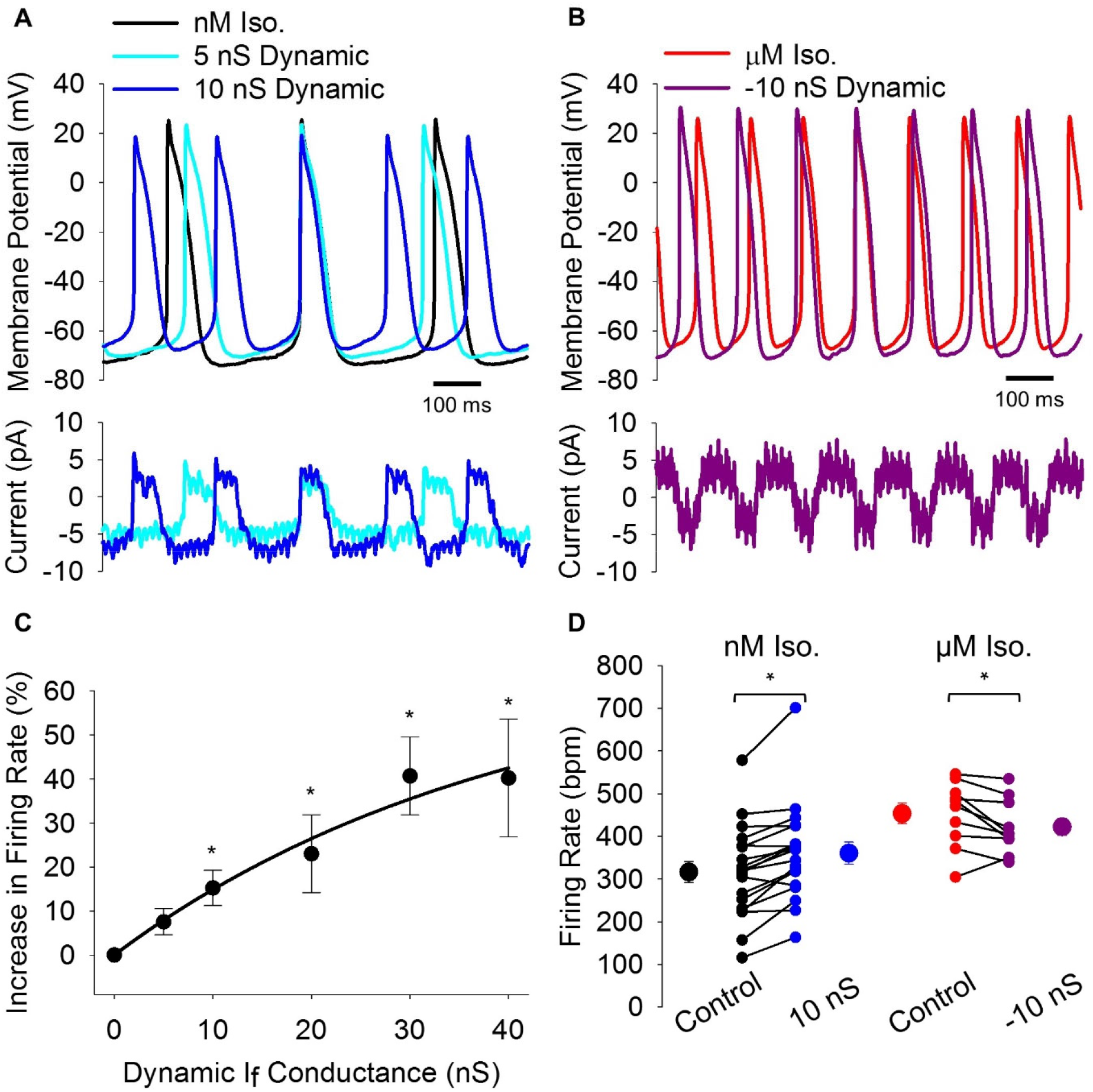
Dynamic Injection of βAR-Stimulated I_f_ Changes AP Firing Rate. **A:** Representative sinoatrial APs (*top*) recorded in 1 nM Iso with no current injection (*black*) and with dynamic addition of βAR stimulated I_f_ (*bottom*) with a conductance of 5 nS (*teal*) or 10 nS (*blue*). **B:** Representative sinoatrial APs recorded in 1 µM Iso (*top*) without current injection (*red*) and with dynamic subtraction of βAR-stimulated I_f_ (*bottom*) with a conductance of 10 nS (*purple*). **C:** Average (±SEM) fractional increase (%) in sinoatrial AP firing rate with dynamic injection of βAR-stimulated I_f_ at conductances between 5 and 40 nS. Injections of 10 nS or greater caused significant acceleration of the AP firing rate compared to control (5 nS - P = 0.9376; 10 nS - P = 0.0401; 20 nS - P = 0.0013; 30 nS - P = 0.0003; 40 nS - P = <0.0001). **D:** Average (±SEM) firing rate of cells perfused with 1 nM (*black*) or 1 µM Iso (*red*) before (*Control*) and after dynamic addition (*blue*) or subtraction (*purple*) of βAR-stimulated I_f_ (10 nS). Individual recordings are shown as smaller circles. Dynamic subtraction of βAR-stimulated I_f_ significantly reduced the AP firing rate compared to 1 µM Iso (P = 0.0402).

In the next set of experiments, we scaled the conductance of the injected current to the endogenous I_f_ measured in each cell to determine cell-specific responses to βAR stimulation of I_f_. In this dynamic clamp protocol, the conductance of the injected βAR-stimulated I_f_ was progressively increased in 2 nS steps (**Fig. S3A**) and the appropriate conductance for each cell was determined *post-hoc* by measuring the endogenous I_f_ during hyperpolarizing pulses following the dynamic clamp recordings (**Fig. S3B**). The endogenous current was measured after the dynamic clamp recordings to allow the use of Ba^2+^ in the isolation of I_f_. Finally, the firing rate and waveform parameters from the scaled dynamic clamp recording were compared to those of APs recorded in 1 nM and 1 µM Iso in the same cell in the absence of current injection (**Fig S3C**).

On average the cell-scaled peak I_f-Iso_ inward current injected during the diastolic depolarization was -9.3 ± 1.1 pA (N = 13) (**Fig. 5A *bottom***), remarkably similar to the 9 pA difference measured between the peak inward I_f_ currents in AP clamp experiments in 1 nM and µM Iso (Peters et al., 2021). Injection of cell-scaled βAR-stimulated I_f_ significantly increased the firing rate by 59 ± 9 AP/min (+22.2% compared to 1 nM Iso; **Table S2**; **Fig. 5**). This increase in firing rate due to βAR stimulation of I_f_ accounted for 41% of the increase in FR seen in response to perfusion of 1 µM Iso (143 ± 24 AP/min; +62.0% compared to 1 nM Iso; **Table S2**; **Fig. 5)**. These data conclusively establish that βAR stimulation of I_f_ contributes significantly to the fight-or-flight increase in AP firing rate in individual SAMs. Importantly, the data also indicate that the full firing rate increase includes contributions from other currents.

**Figure 5:**
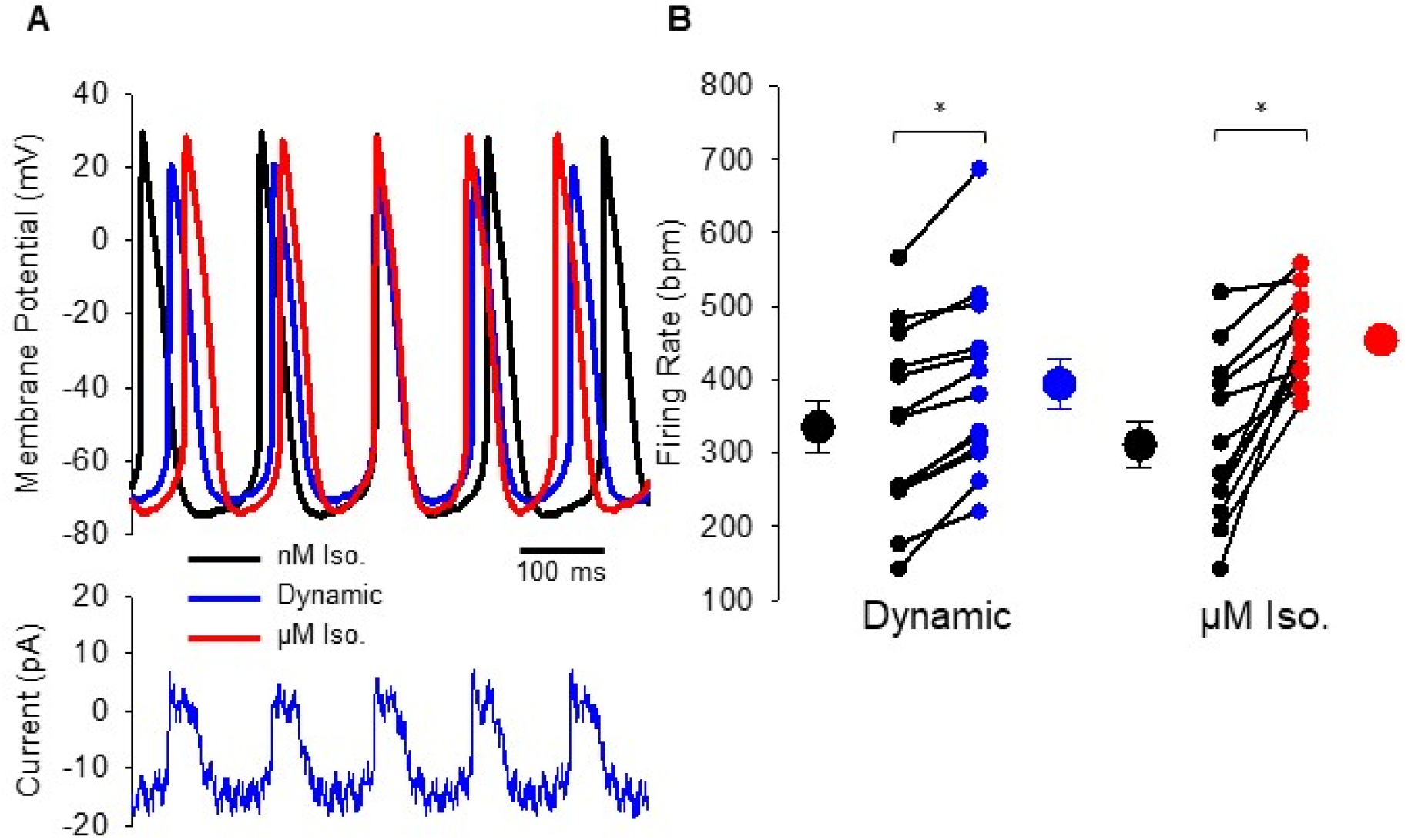
Cell-scaled dynamic clamp shows βAR Stimulation of I_f_ accounts for 36% of the increase in AP firing rate. **A *top*:** Representative sinoatrial APs from one cell in 1 nM Iso with no current injection (*black*), in 1 nM Iso with dynamic injection of βAR-stimulated I_f_ (*blue*), and in 1 µM Iso with no current injection (*red*). **A *bottom*:** Dynamic current injection (*blue*) during the APs recorded during dynamic clamp. **B:** Average (± SEM) firing rate of cells in 1 nM Iso before (*black*) and after cell-scaled dynamic injection of βAR stimulated I_f_ (*blue*) or perfusion of 1 µM Iso (*red*). Individual recordings are shown as small circles. Control AP firing rates immediately preceding the dynamic I_f-Iso_ addition and perfusion of 1 µM Iso did not differ significantly (P = 0.7432).

We next analyzed the effects of dynamic injection of I_f-Iso_ on the AP waveform parameters identified by correlation analysis and machine learning as the primary drivers of the firing rate increase with βAR stimulation, namely EDD, LDD, DDR, and LAPD. We found that I_f-Iso_ accounted for 80% of the decrease in LAPD, 50% of the decrease in DD, and 40% of the decrease in LDD seen with perfusion of 1 µM Iso (**Table S2**; **Fig. 6A** and **B**). While dynamic injection of I_f-Iso_ elicited trends towards faster DDR and shorter EDD, these effects were not statistically significant and the effects in individual cells were variable. Ultimately, these data suggest that βAR stimulation of I_f_ contributes to the fight-or-flight heart rate increase through shortening of the diastolic depolarization and late repolarization phases of the sinoatrial AP resulting in a 60% reduction in the time between APs.

**Figure 6:**
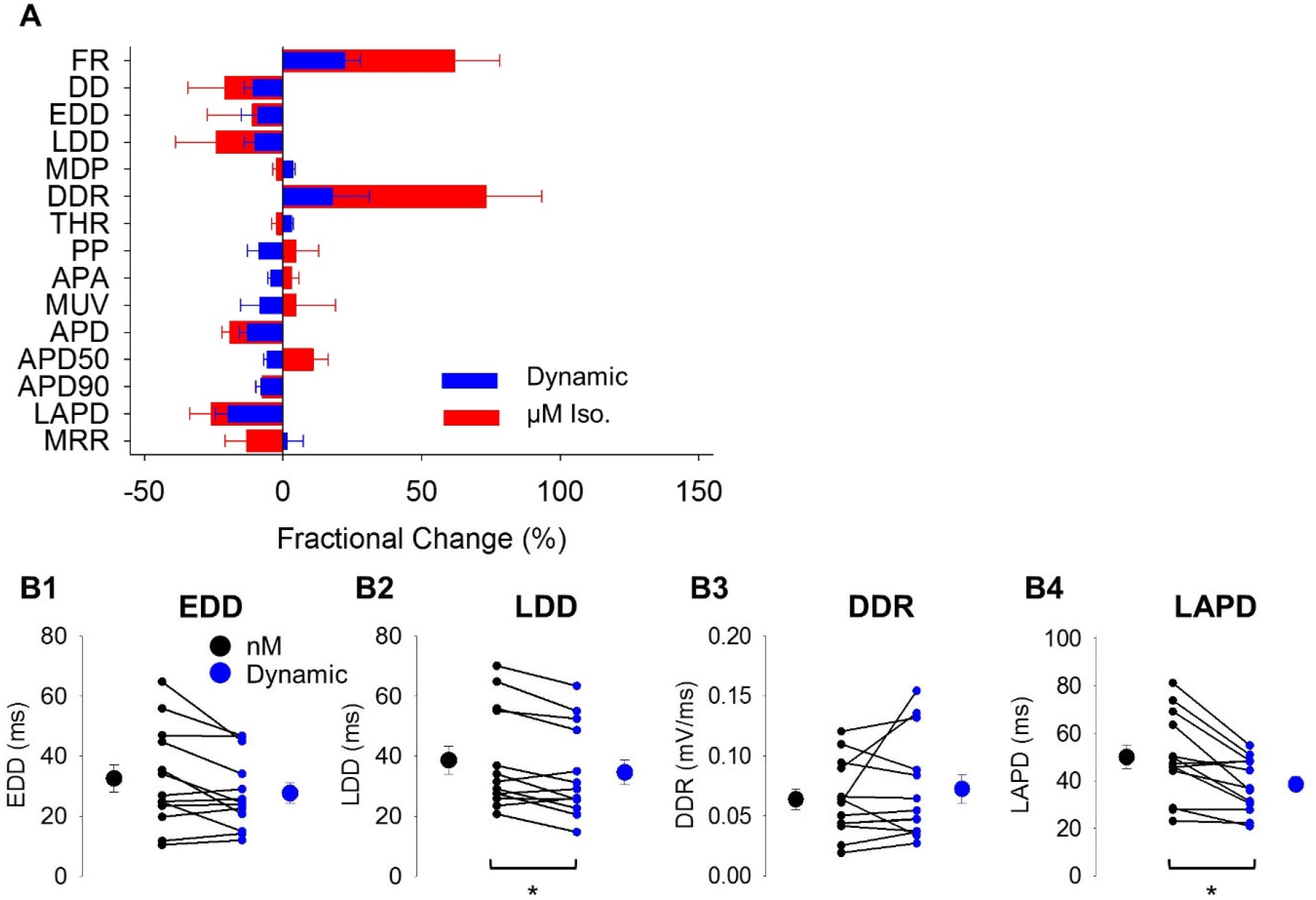
βAR Stimulation of I_f_ Significantly Shortens LAPD and DD. **A:** Average (±SE) fractional change (%) in sinoatrial AP firing rate and waveform parameters with perfusion of 1 µM Iso (*red*) or dynamic injection of βAR stimulated I_f_ (*blue*) compared to recordings in 1 nM Iso. **B1-B4:** Average (+SEM) waveform parameters (EDD, LDD, DDR, LAPD) of cells in 1 nM Iso before (*black*) and after cell-scaled dynamic injection of βAR stimulated I_f_ (*blue*). Individual recordings are shown as small circles. Details of statistical tests can be found in **Supplementary Table S2**.

## Discussion

The sympathetic nervous system fight-or-flight increase in heart rate is among the most fundamental of physiological responses in mammals. Yet despite decades of research, the mechanisms underlying the increase in spontaneous AP firing rate in SAMs remain unresolved (Lakatta and DiFrancesco, 2009; Bucchi et al., 2012; Hennis et al., 2021). In this study, we first show that while virtually all aspects of the sinoatrial AP are changed by βAR stimulation, the magnitude of the fight-or-flight increase in firing rate is mediated by changes in only a subset of AP waveform parameters. Importantly, this subset of parameters constrains the list of βAR sensitive processes that contribute to the fight-or-flight response. We then quantitatively establish that I_f_ accounts for about 41% of the fight-or-flight increase in AP firing rate in SAMs, countering previous claims that I_f_ does not play a role in this process.

### βAR stimulation increases firing rate via a subset of AP waveform parameters

Numerous studies have suggested that βAR stimulation increases AP firing rate at least in part by increasing the DDR in SAMs (Zhang and Vassalle, 2003; Bucchi et al., 2007; Verkerk et al., 2012; Larson et al., 2013; Protze et al., 2017). However, more definitive determination of the sinoatrial AP waveform changes that mediate the fight-or-flight increase in firing rate have been hampered by inconsistent parameter definitions and small datasets. Our use of automated AP waveform analysis of a large dataset provides the necessary resolution to conclusively show that βAR stimulation significantly alters almost all AP waveform parameters — consistent with the host of βAR-sensitive processes in SAMs.

Importantly, however, our correlation analysis and random-forest computer learning (**Fig. 3**) constrain the list of βAR-sensitive processes involved in the fight-or-flight response by showing that a subset of sub-threshold parameters — namely EDD, LDD, DDR, and LAPD — are the key determinants of AP firing rate and the βAR-mediated increase in firing rate in SAMs. This finding quantitatively confirms prior suggestions that shortening of the interval between APs is the main driver of autonomic regulation of heart rate and that the currents active during these phases of the AP are the key mechanisms for the fight-or-flight increase in heart rate (Bucchi et al., 2007; DiFrancesco, 2010). In contrast, changes in systolic parameters such as shortening of the APD90 only account for 10-20% of the total change in cycle length and differ little between cells with small and large βAR-stimulated firing rate increases (**Fig. 3C2**).

### Evaluation the role of I_f_ in the fight-or-flight response is complicated by reduced resting rates

Whether and to what degree I_f_ is responsible for cardiac pacemaking and autonomic regulation of heart rate has long been debated (Lakatta and DiFrancesco, 2009; Hennis et al., 2021). On one hand, the slow rate of activation and hyperpolarized voltage-dependence of I_f_ seemingly preclude appreciable activation of I_f_ during the sinoatrial AP, even when βARs are stimulated (Lakatta and DiFrancesco, 2009). On the other hand, an essential role for I_f_ in cardiac pacemaking is clearly indicated by slower heart rates and sinus arrhythmias in response to I_f_ blockers and HCN4 mutations (Stieber et al., 2003; Herrmann et al., 2007; Hoesl et al., 2008; Harzheim et al., 2008; Alig et al., 2009; Baruscotti et al., 2011; Bucchi et al., 2012; Mesirca et al., 2014; Fenske et al., 2020). Our recent work using the AP-clamp technique helps to resolve this paradox by showing that I_f_ is constitutively active throughout the duration of the sinoatrial AP. This persistent activation allows I_f_ to conduct more than 50% of the net inward charge movement during the AP under control conditions, despite its low fractional open probability (Peters et al., 2021). Importantly, our study also demonstrated that βAR stimulation increases the fraction of net inward charge movement conducted by I_f_ to 93% of the total (Peters et al., 2021) providing strong support for a substantive role for I_f_ in the fight-or-flight response in SAMs.

A role for I_f_ in the βAR-mediated increase in sinoatrial AP firing rate has been suggested since the earliest description of I_f_ showed that adrenaline potentiates the current by shifting the activation voltage dependence to more positive potentials (Brown et al., 1979). Yet little direct data exists to support a role for I_f_ in the increase in AP firing rate, owing in part to the difficulty of isolating the effects of a single current.

A number of studies through the years have used I_f_ blockers and mice bearing mutations in HCN4 to probe the role of I_f_ in the fight-or-flight response. These manipulations consistently reduce resting and maximal heart rates in animals and spontaneous AP firing rates in isolated SAMs but do not prevent a partial increase in pacemaker activity from the slower baseline rates (Leitch et al., 1993; Cai et al., 1995; Du et al., 2004; Stieber et al., 2006; Bucchi et al., 2007; Swedberg et al., 2010; Sosunov and Anyukhovsky, 2012). Interpretation of such results is problematic for a number of reasons. For example, I_f_ blockers such as Cs^+^ and ivabradine can cause incomplete block at commonly used concentrations (DiFrancesco, 1982; Bucchi et al., 2002; Peters et al., 2021) and/or non-specific effects on other currents (Lees-Miller et al., 2003; Haechl et al., 2019; Peters et al., 2021), both of which confound estimates of the role of I_f_.

Meanwhile, knockout of HCN4 or disruption of cAMP binding are embryonically lethal (Stieber et al., 2003; Harzheim et al., 2008), there are differences in phenotype between global and cardiac-specific conditional knockouts (Herrmann et al., 2007; Hoesl et al., 2008; Baruscotti et al., 2011), and models that reduce the cAMP sensitivity of HCN4 either truncate large portion of the channel or do not disrupt PKA-dependent phosphorylation of the channel, an often overlooked activator of I_f_ during βAR stimulation (Liao et al., 2010).

It has been suggested that βAR-mediated increases in heart rate or firing rate from the lower baseline levels in these models exclude a role for I_f_ in the fight-or-flight response (Zhang and Vassalle, 2003; Lakatta and DiFrancesco, 2009; Alig et al., 2009; Mesirca et al., 2014; Fenske et al., 2020; Hennis et al., 2021). However, this argument assumes that firing rate acceleration in a cell firing 200 AP/min and a cell firing 400 AP/min require a similar amount of depolarizing current. This is clearly not the case: the shorter diastolic interval in faster cells requires a larger depolarizing current to affect the same change in membrane potential to reach threshold. Thus, in slower cells — such as when I_f_ is reduced by blockers or mutations — βAR stimulation may have sufficient time to produce a “normal” increase in firing rate via additional pathways, whereas in faster cells the same incomplete stimulation would produce a smaller increase in firing rate. Indeed, analysis of all the APs recorded in the present study show that a larger βAR-mediated increase in AP firing rate is correlated with a larger increase in DDR (**Fig. 3A4**). Moreover, analysis of cells binned by firing rate reveals that a 118 AP/min increase in mean firing rate in faster cells (from 338 to 456 AP/min, N=54) requires that the diastolic depolarization rate increase by 0.044 mV/ms, which is about three times the DDR increase required for a similar magnitude firing rate increase (121 AP/min) in slower cells (217 to 338 AP/min, N=54; corresponding to a DDR increase of 0.015 mV/ms; **Table S3**).

Taken together, previous studies consistently demonstrate a residual fight-or-flight increase in pacemaking in the absence of I_f_, clearly indicating that I_f_ alone is insufficient to produce the full response. However, prior studies do not preclude a contribution of I_f_ to the response given the reductions in basal rates in all of the models.

### βAR stimulation of I_f_ alone increases AP firing rate through changes to the diastolic depolarization

To quantitatively determine the role that βAR stimulation of I_f_ plays in firing rate acceleration in SAMs, we used the dynamic clamp technique with AP-clamp validated models parameterized to I_f_ currents recorded from SAMs under identical conditions (Peters et al., 2021). The dynamic clamp currents in our experiments were scaled to the endogenous I_f_ directly measured in each cell (**Figure S2**). While a prior study used dynamic clamp in rabbit SAMs to investigate the role of βAR stimulation of I_f_ in that system, the I_f_ conductance was scaled indirectly, via a calibration curve constructed by relating I_f_ amplitude to firing rate in a mathematical model of the sinoatrial AP (Severi et al., 2012; Ravagli et al., 2016). Unfortunately, this scaling paradigm is recursive, as it assumes that differences in firing rate are exclusively due to changes in the I_f_ amplitude. Moreover, the I_f_ models used in the prior study only accounted for βAR-stimulated shifts in the voltage-dependence I_f_ but not the changes in kinetics, despite the fact that βAR speeding of activation and slowing of deactivation of I_f_ are fundamental to its role in pacemaking (Peters et al., 2021). Despite these differences in approach and system, the results of the two dynamic clamp studies are comparable, with our data showing that βAR-stimulated I_f_ drives 41% of the increase in AP firing rate in mouse SAMs (**Fig. 5**) similar to the ∼ 38% fraction in rabbit SAMs (Ravagli et al., 2016).

Our study further leveraged automated AP waveform analysis to additionally demonstrate that βAR modulation of I_f_ accounts for 60% of the shortening between APs, including 50% of the DD shortening and 80% of the LAPD shortening that occur with βAR stimulation (**Fig. 6**). Since these parameters are primary determinants of the βAR increase in AP firing rate determined by correlation analysis and machine learning (**Fig. 3**), they further high the importance of I_f_ in the fight-or-flight response in SAMs. Although the LAPD — defined as the time from when the threshold potential is reached during the repolarization until the subsequent maximum diastolic potential (**Table 1; Fig. S1**) — accounts for nearly 25% of the total cycle length and a third of the time spent in diastole, we are unaware of any prior studies that quantified its duration. The prominent role for I_f_ during this phase of the AP is consistent with our recent finding that I_f_ remains active throughout the AP (Peters et al., 2021) and the βAR- stimulated increase in I_f_ is uniquely well-suited to act on LAPD when other depolarizing currents are not predicted to be active.

Equally important to the AP parameters that change upon I_f_ injection are those that do not change. Compared to isoproterenol stimulation, injection of βAR-stimulated component of I_f_ alone produced opposite effects on the MDP, THR, PP, APA, MUV, APD_50_, and MRR (**Fig. 6A**). The opposite effects of isoproterenol and I_f-Iso_ on the MDP indicate that sympathetic stimulation must also increase an outward current that is active during the diastolic depolarization, possibly a voltage-gated or calcium-sensitive SK or BK potassium channel (Torrente et al., 2017; Lai et al., 2014). Similarly, isoproterenol must stimulate an inward current — such as Na_v_1.1 or Ca_v_1.2 (Hagiwara et al., 1988; Lei et al., 2004) — during the AP upstroke to depolarize the APA and the MUV. Furthermore, it is clear that inward current through NCX also contributes to the fight-or-flight response and further shortening of DD (Lakatta et al., 2010; Gao et al., 2013).

### Conclusions

While our direct measurements provide a uniquely quantitative resolution of the contribution of I_f_ to the fight-or-flight response, they also clearly indicate that other currents are also required. Given the fundamental importance of the fight-or-flight increase in heart rate in mammals, it is reassuring to know that redundant mechanisms are involved. As noted by Lipsius and Bers nearly two decades ago, “it is … prudent to adopt a more balanced integrative view of the cardiac pacemaker, and acknowledge the real interplay of many factors” (Lipsius and Bers, 2003). The challenge for future studies is to resolve the coordinated and interrelated responses of multiple molecular processes.

## Materials and Methods

### Animal Care and Euthanasia

This study was carried out in accordance with the US Animal Welfare Act and the National Research Council’s *Guide for the Care and Use of Laboratory Animals* and was conducted according to a protocol that was approved by the University of Colorado-Anschutz Medical Campus Institutional Animal Care and Use Committee. Six-to eight-week old male C57BL/6J mice were obtained from Jackson Laboratories (Bar Harbor, ME, USA). Animals were anesthetized by isofluorane inhalation and euthanized under anesthesia by cervical dislocation and bilateral thoracotomy.

### Acute Isolation of Murine Sinoatrial Myocytes

Sinoatrial myocytes were isolated as we have previously described (Sharpe et al., 2016; Larson et al., 2013). The heart was excised and the sinoatrial node dissected in Tyrode’s solution (in mM: 140 NaCl, 5.4 KCl, 5.6 Glucose, 5 HEPES, 1.8 CaCl_2_, 1.2 KH_2_PO_4_, 1 MgCl_2_; Ph adjusted to 7.4 with NaOH) containing 10 USP units/ml heparin (Sagent Pharmaceuticals, Schaumburg, IL, USA) at 36°C. Strips of sinoatrial node tissue were subsequently rinsed in a low Ca^2+^ modified Tyrode’s solution (in mM: 140 NaCl, 50 Taurine, 5.4 mM KCl, 18.5 Glucose, 5 HEPES, 1.2 KH_2_PO_4_, 0.066 CaCl_2_, 1 mg/ml BSA; pH adjusted to 6.9 with NaOH) at 36°C. The tissue was digested for 25-30 min in 2 ml of the same solution containing 1.79 U protease type XIV (Sigma-Aldrich, St. Louis, MO, USA), 0.54 U collagenase B (Sigma-Aldrich, St. Louis, MO, USA) or 1064 U type II collagenase (Worthington), 9 U elastase (Worthington), and 652 µg type XIV protease (Sigma) for 30 min. After enzymatic digestion, the tissue was extensively rinsed in a modified Kraft-Brühe (KB) solution (in mM: 100 K-glutamate, 25 KCl, 20 glucose, 20 taurine, 10 K-aspartate, 10 KH_2_PO_4_, 5 creatine, 5 HEPES, 2 MgSO_4_, 0.5 EGTA, 1 mg/ml BSA; pH adjusted to 7.2 with KOH) at 36°C. Individual sinoatrial myocytes were obtained by mechanical trituration of the digested tissue with a fire-polished glass pipette for 2- 5 min in 2.5 ml of the KB solution. The calcium concentration of the cell suspension was gradually increased to a final concentration of 1.8 mM during an adaptation time of 30 min. Cells were stored at room temperature for up to 5 hours before recordings.

### Electrophysiology

Aliquots of cells (∼100 µl) were transferred to a recording chamber on the stage of an inverted microscope and cells were continuously perfused at 1-2 ml/min with Tyrode’s solution with or without the β adrenergic receptor agonist, isoproterenol (Iso; 1 nM to 10 µM) at a temperature of 35 ± 2°C. Data were acquired at 50 kHz and low-pass filtered at 10 kHz using an Axopatch 200B amplifier, Digidata 1440a A/D converter, and ClampEx software (Molecular Devices). Bath and perfusing solutions were maintained at 35ºC for all recordings with a feedback-controlled temperature controller (TC-344B, Warner Instruments). Recording pipettes were pulled from borosilicate glass (VWR International) using a Sutter Instruments P-97 horizontal puller. Pipette resistances ranged from 1.5 to 5 MΩ.

Amphotericin B perforated patch clamp recordings were performed as previously described (Sharpe et al., 2017). The perforated patch pipette solution contained (in mM): 135 KCl, 0.1 CaCl_2_, 1 MgCl_2_, 5 NaCl, 10 EGTA, 4 MgATP, 10 HEPES; adjusted to pH 7.2 with KOH. A perforation stock solution was prepared by solubilizing 50 mg/ml Pluronic F-127 in DMSO (both from Sigma-Aldrich, St. Louis, MO, USA) and adding 20 mg/ml Amphotericin-B (Apexbio Technology LLC, Houston, TX, USA). 5-15 µl of the perforation stock solution was then added to 500 µl of the pipette solution and solubilized by sonication for a final Amphotericin-B concentration of 0.2-0.6 mg/ml.

After establishing a high-resistance seal (>1 GΩ), the perforation progress was confirmed by a reduction in access resistance in voltage-clamp mode and by a visible minimization of MDP to a stable value in current-clamp mode. Access resistance values were usually in the range of 10 ± 2 MΩ, while MDPs regularly stabilized around -69 mV after roughly 30 s of perforation time (Table S1).

Spontaneous APs were recorded in current-clamp mode in the initial condition until both MDP and FR were stable for at least 1 min, usually within 1-3 min after establishing a high-resistance seal. The perfusing solution was then changed to a different Iso concentration. In general, changes in FR were completed within 30 s after the solution change and the recording was then continued for at least another 1 min to document the Iso-sensitive changes of AP waveform parameters.

### Dynamic Clamp Experiments

For dynamic clamp experiments, we modified an Arduino-based system (Desai, 2017) to better match the system’s digital output range with the current command input range of the Axopatch 200B patch clamp amplifier. Control and feedback from the system were measured using the custom open-source “pyClamp” (Rickert and Proenza, 2019a) user client and “dyClamp” (Rickert and Proenza, 2019b) host controller software sketches.

Dynamic current-clamp recordings from SAMs were performed in the perforated-patch clamp configuration. SAMs were continuously perfused with Tyrode’s solution containing either 1 nM or 1 µM Iso until both FR and MDP were stable for approximately 1 min. Dynamically-calculated currents were then injected into the SAMs based on I_f_ equations from previously described and validated current models for I_f_ in both 1 nM and 1 µM Iso (**Fig. S2A**) (Peters et al., 2021). Both models are Hodgkin-Huxley type models with fast and slow activation and deactivation time constants to account for the complex I_f_ kinetics. The 1 µM Iso model accounts for the depolarizing shift in the steady-state activation, slower deactivation, and faster channel activation (Peters et al., 2021).

The Iso difference current was calculated as the difference between the 1 nM and 1 µM models at each time step. The Iso difference current was added to cells perfused with 1 nM Iso to simulate the effects of an increase in Iso concentration, while the Iso difference current was subtracted from cells perfused in 1 µM Iso to simulate the effects of reducing the Iso concentration. In each cell, following a control period of 30 s of APs in the initial Iso concentration, the Iso difference current was injected in a series of steps of increasing maximum conductance in the range of 5-40 nS (corresponding to current injections of approximately 1-25 pA). Dynamic current injections were monitored with both the current command input to the amplifier and the 10 kHz-filtered head stage signal returned from the amplifier. For each conductance level, currents were injected for approximately 1 minute to allow both MDP and/or FR to stabilize. The stable APs recorded during the latter part of the current injection time were then analyzed.

To measure the cell-specific effects of βAR stimulation of I_f_, cells were patched in the perforated-patch configuration and continuously perfused with 1 nM Iso. Baseline AP firing was measured in current clamp mode with zero injected current until both FR and MDP were stable (typically within 1 min). Dynamically calculated currents were then injected into the SAMs in steps of 2 nS for 15 s each up to 20 nS as described above (**Fig. S2B**). Finally, the dynamic current injection was removed and 1 µM Iso was perfused to measure the full cell-specific βAR response. Following AP recordings in 1 µM Iso, the amplifier was switched to voltage-clamp mode and whole-cell access was gained to measure cellular capacitance. 1 mM BaCl_2_ was perfused on to the cell and inward I_f_ was then measured as the activating current during a 3 s pulse to -130 mV in voltage-clamp mode (**Fig. S2C**). The cell specific maximal I_f_ conductance (G_f_) was calculated using equation 1:

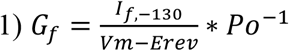

Where I_f,-130_ is the amplitude of the activating inward current at -130 mV, V_m_ is the membrane potential (−130 mV), E_rev_ is the reversal potential of I_f_ (−30 mV), and P_o_ is the predicted steady-state open probability of the 1 µM Iso model at -130 mV (∼100%). The FR and AP waveform during the dynamic injection with a conductance closest to that measured in the cell (within 1 nS) was compared to that measured in the same cell in 1 nM (**Fig. S2D**). The difference in FR and AP waveform with and without dynamic injection was then compared to difference between 1 nM and 1 µM Iso in the same cell. Because the firing rate runs-down slightly during the protocol (typically <20 AP/min over 10 min), the temporally closest 1 nM recording was used for comparison to each of dynamic injection and βAR stimulation.

### Action Potential Waveform Analysis with ParamAP

AP waveforms were parameterized using the ParamAP software (Rickert and Proenza, 2017, 2021). Segments with APs were extracted from the recordings and saved in Axon’s text file format (ATF) before being analyzed with ParamAP. Variable length analysis windows were used to capture APs with stable features and membrane parameters and to avoid noise artifacts. ParamAP was updated to include the late-action potential duration (LAPD), which measures the time in diastole preceding the maximum diastolic potential. All parameter definitions are listed in Table 1. Importantly, the measured waveform parameters captured by ParamAP from individual cells can be used to recapitulate sinoatrial APs with linear segments defined by said parameters (**Fig. S1B**). This indicates that ParamAP is not under-parameterizing the sinoatrial AP.

### Machine Learning

We used machine learning to predict the changes in SAM firing rate given measured changes in AP waveform parameters (**Table 1**). We constructed 100 random forest models, each populated with 1000 trees, using the scikit-learn Python module (Pedregosa et al., 2011). Models between 50 and 5000 trees were tested and similar results obtained for each model, with 1000 and 5000 tree models showing less variability between any given forest. Each of the random forests was generated with a pseudo-random starting point to generate a range of feature importance measures. Models were trained on the dataset of 50 cells recorded in 1 nM and 1 µM Iso to predict the fractional change in firing rate with βAR stimulation. The predicting variables were chosen to measure all relevant periods, membrane potentials, and rates, while excluding variables that measured subsets of the AP waveform already included within another predictor. For example, APD50 was not included as that region of the AP-waveform is a subset of the data included in the APD90 variable. Models were tested against the effects of βAR stimulation in the independent dataset of cells used for dynamic clamp analysis and were capable of predicting the change in firing rate with an average error of 12.2% (approximately the standard error of the mean for the firing rate change in the training and targets datasets, 13.4% and 11.0%, respectively). Predictor importance was measured by evaluating the contribution of each parameter to reducing node impurity within the training data set using the default function within the scikit-learn module. Thus, importance is both a measure of the frequency with which a predictor appears within the forest and its ability to split data into less variable subgroups.

### Statistical Analysis

All statistical analysis was performed in JMP 15 (SAS Institute). Comparisons of unmatched 0 nM, 1 nM, 1 µM, and 10 µM Iso were performed using a 1-way ANOVA. Pairwise comparisons were performed using a TUKEY Post-hoc test. Comparisons of matched 1 nM and 1 µM Iso datasets were performed using paired t-tests. Comparisons in dynamic clamp experiments were performed using paired t-tests with a Bonferroni correction of 2 used for multiple pairwise comparisons (1 nM Iso vs. dynamic and 1 µM-1 nM Iso vs dynamic-1 nM Iso). Correlations between parameters were calculated using a Pearson correlation coefficient. Statistical significance was set at P < 0.05.

## Abbreviation

βAR: β-adrenergic receptor
AP: Action potential
APA: Action potential amplitude
APD: Action potential duration
APD_50_: Action potential duration at 50% repolarization
APD_90_: Action potential duration at 90% repolarization
cAMP: Cyclic adenosine monophosphate
CL: Cycle length
DD: Diastolic duration
DDR: Diastolic depolarization rate
EAPD: Early AP duration
EDD: Early diastolic duration
FR: Firing rate
HCN: Hyperpolarization-activated cyclic nucleotide-sensitive channel
HR: Heart rate
Iso: Isoproterenol
LAPD: Late AP duration
LDD: Late diastolic duration
MDP: Maximum diastolic potential
MRR: Maximum repolarization rate
MUV: Maximum upstroke velocity
PP: Peak potential
SAM: Sinoatrial node myocyte
THR: Threshold potential

## Acknowledgements

This work was funded by NIH grants R01 HL088427 to C.P., R01HL131517, R01HL141214, and P01HL141084; the Stimulating Peripheral Activity to Relieve Conditions Grants OT2OD026580 to E.G. and R00HL138160 to S.M.; and by postdoctoral fellowships from the American Heart Association (19POST34380777 and 830889) to C.H.P. The Optogenetics and Neural Engineering (ONE) Core at the University of Colorado School of Medicine provided engineering support for this research. The ONE Core is part of the NeuroTechnology Center, funded in part by the School of Medicine and by the National Institute of Neurological Disorders and Stroke of the National Institutes of Health under award number P30NS048154. Dr. Debashis Ghosh (University of Colorado, Anschutz Medical Campus, Department of Biostatistics and Informatics) provided consulting on machine learning analysis for this paper.

## Supplementary Materials

**Table S1:**
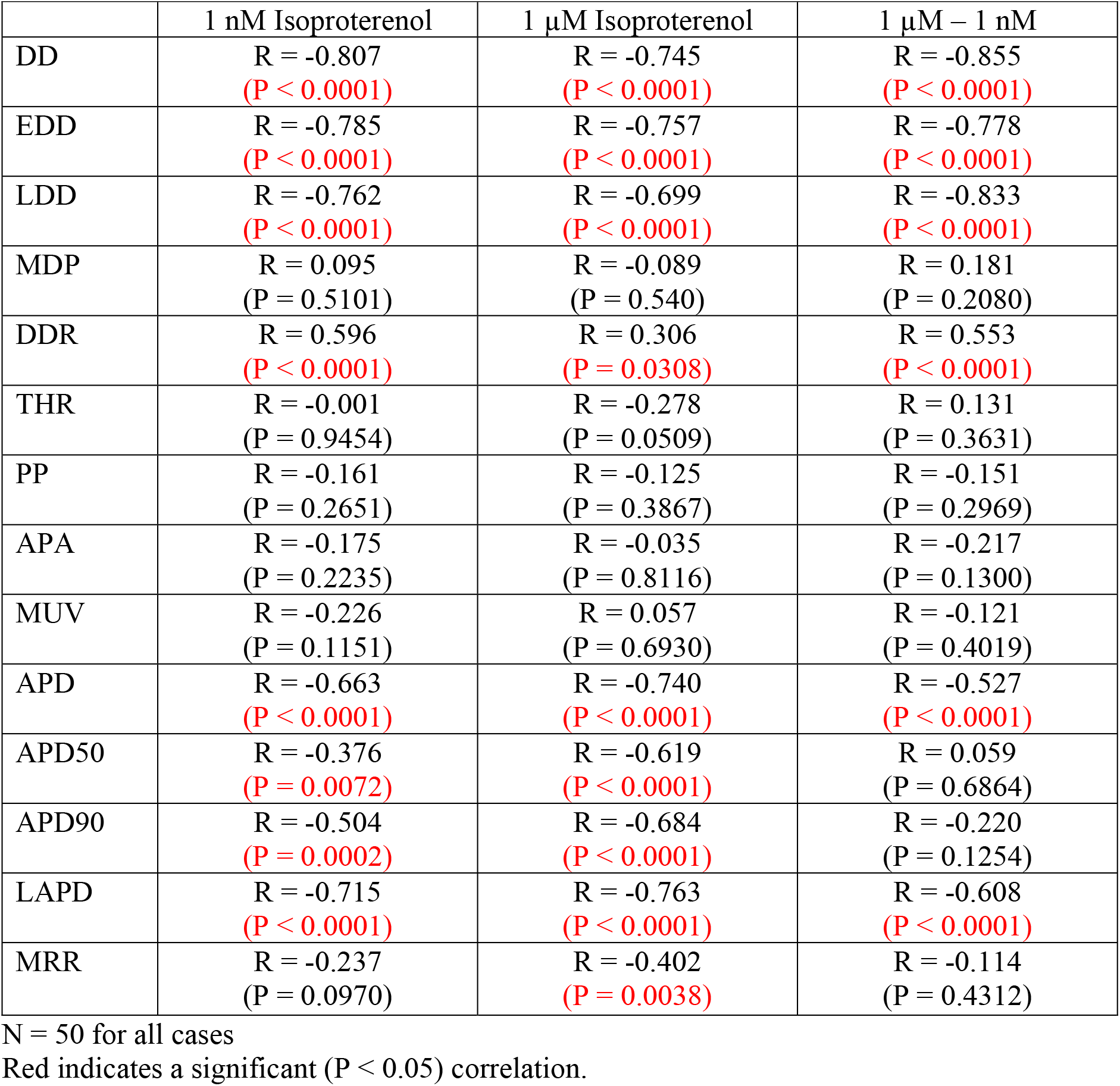
Pearson Correlation Coefficients Between Firing Rate and Waveform Parameters in 1 nM and 1 µM Isoproterenol.

| | 1 nM Isoproterenol | 1 μM Isoproterenol | 1 μM – 1 nM |
| --- | --- | --- | --- |
| DD | R = -0.807<br>(P < 0.0001) | R = -0.745<br>(P < 0.0001) | R = -0.855<br>(P < 0.0001) |
| EDD | R = -0.785<br>(P < 0.0001) | R = -0.757<br>(P < 0.0001) | R = -0.778<br>(P < 0.0001) |
| LDD | R = -0.762<br>(P < 0.0001) | R = -0.699<br>(P < 0.0001) | R = -0.833<br>(P < 0.0001) |
| MDP | R = 0.095<br>(P = 0.5101) | R = -0.089<br>(P = 0.540) | R = 0.181<br>(P = 0.2080) |
| DDR | R = 0.596<br>(P < 0.0001) | R = 0.306<br>(P = 0.0308) | R = 0.553<br>(P < 0.0001) |
| THR | R = -0.001<br>(P = 0.9454) | R = -0.278<br>(P = 0.0509) | R = 0.131<br>(P = 0.3631) |
| PP | R = -0.161<br>(P = 0.2651) | R = -0.125<br>(P = 0.3867) | R = -0.151<br>(P = 0.2969) |
| APA | R = -0.175<br>(P = 0.2235) | R = -0.035<br>(P = 0.8116) | R = -0.217<br>(P = 0.1300) |
| MUV | R = -0.226<br>(P = 0.1151) | R = 0.057<br>(P = 0.6930) | R = -0.121<br>(P = 0.4019) |
| APD | R = -0.663<br>(P < 0.0001) | R = -0.740<br>(P < 0.0001) | R = -0.527<br>(P < 0.0001) |
| APD50 | R = -0.376<br>(P = 0.0072) | R = -0.619<br>(P < 0.0001) | R = 0.059<br>(P = 0.6864) |
| APD90 | R = -0.504<br>(P = 0.0002) | R = -0.684<br>(P < 0.0001) | R = -0.220<br>(P = 0.1254) |
| LAPD | R = -0.715<br>(P < 0.0001) | R = -0.763<br>(P < 0.0001) | R = -0.608<br>(P < 0.0001) |
| MRR | R = -0.237<br>(P = 0.0970) | R = -0.402<br>(P = 0.0038) | R = -0.114<br>(P = 0.4312) |

N = 50 for all cases
Red indicates a significant (P < 0.05) correlation.

**Table S2:**
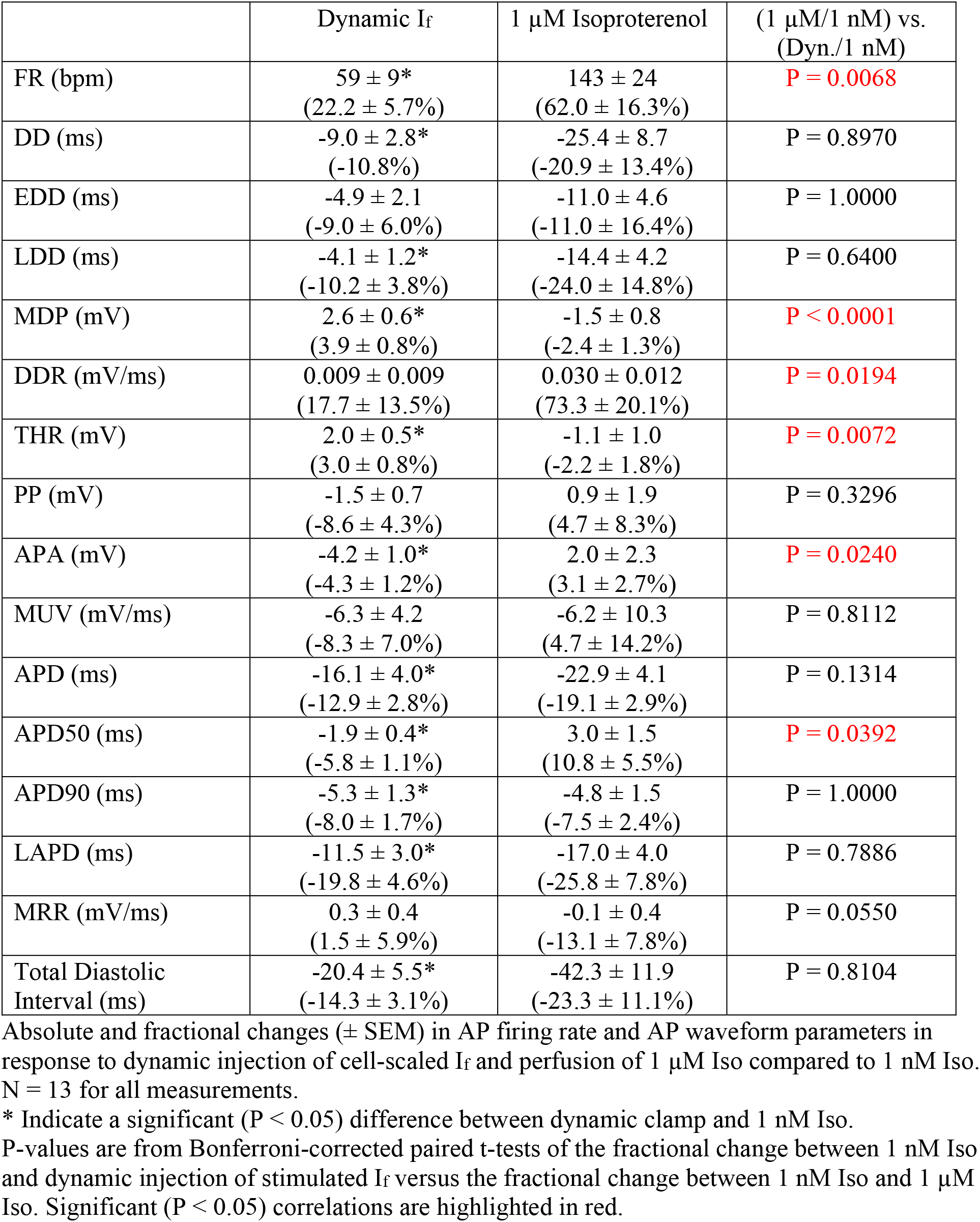
Dynamic Clamp Changes in AP firing rate and AP waveform.

**Table S3:**
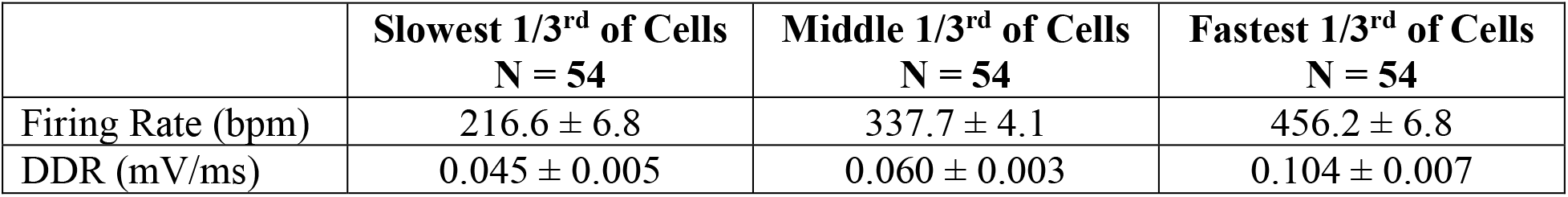
Diastolic Depolarization Rate in Cells Binned by Firing Rate.

**Figure S1:**
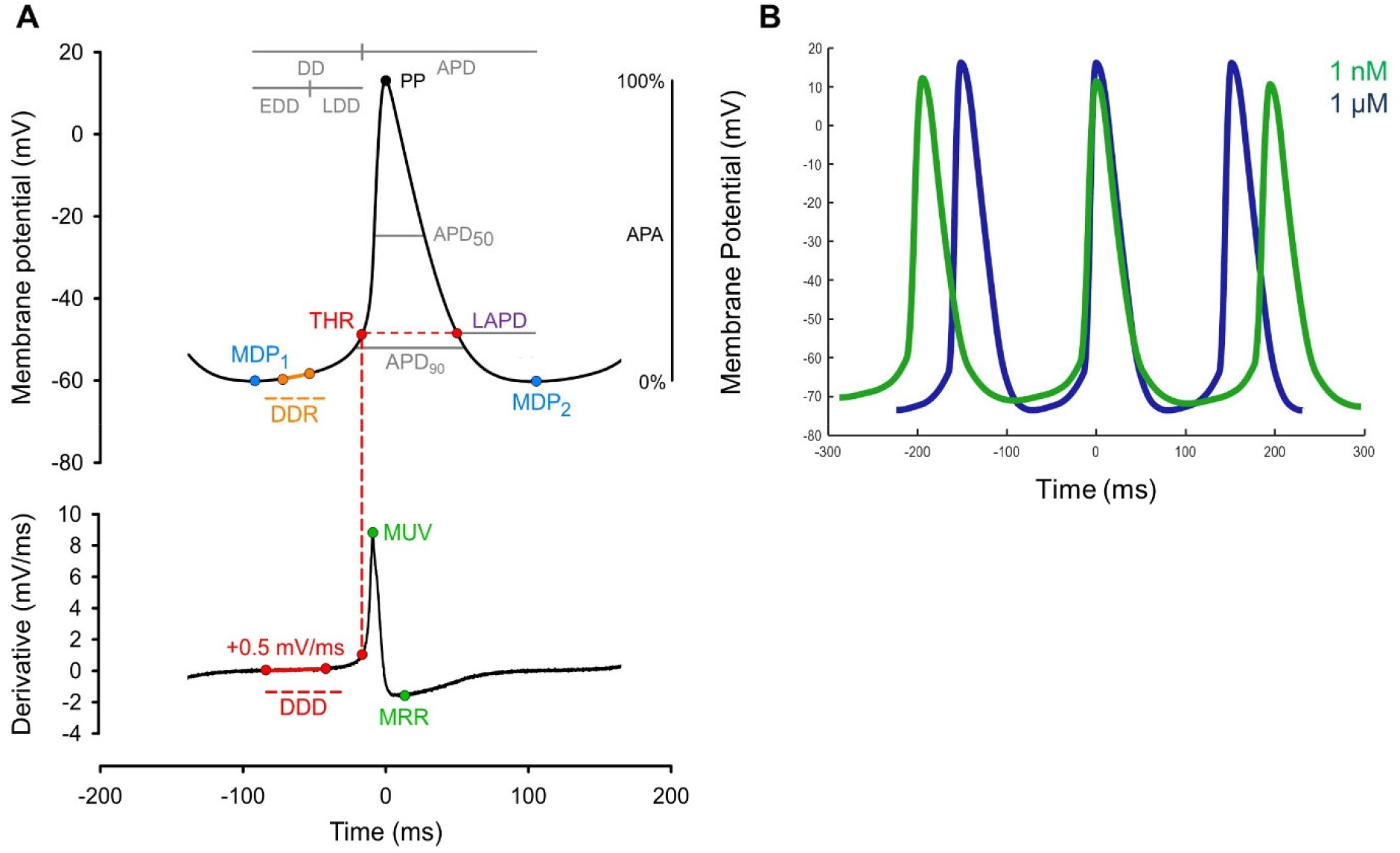
ParamAP Automates the Capture of Sinoatrial Node AP Waveform Parameters. **A:** Schematic illustration of ParamAP waveform parameters, including LAPD (*purple*). **B:** Sinoatrial AP waveforms modeled by line segments representing average waveform parameters determined by ParamAP in 1 nM or 1 µM Iso indicate that ParamAP adequately parameterizes the sinoatrial AP.

**Figure S2:**
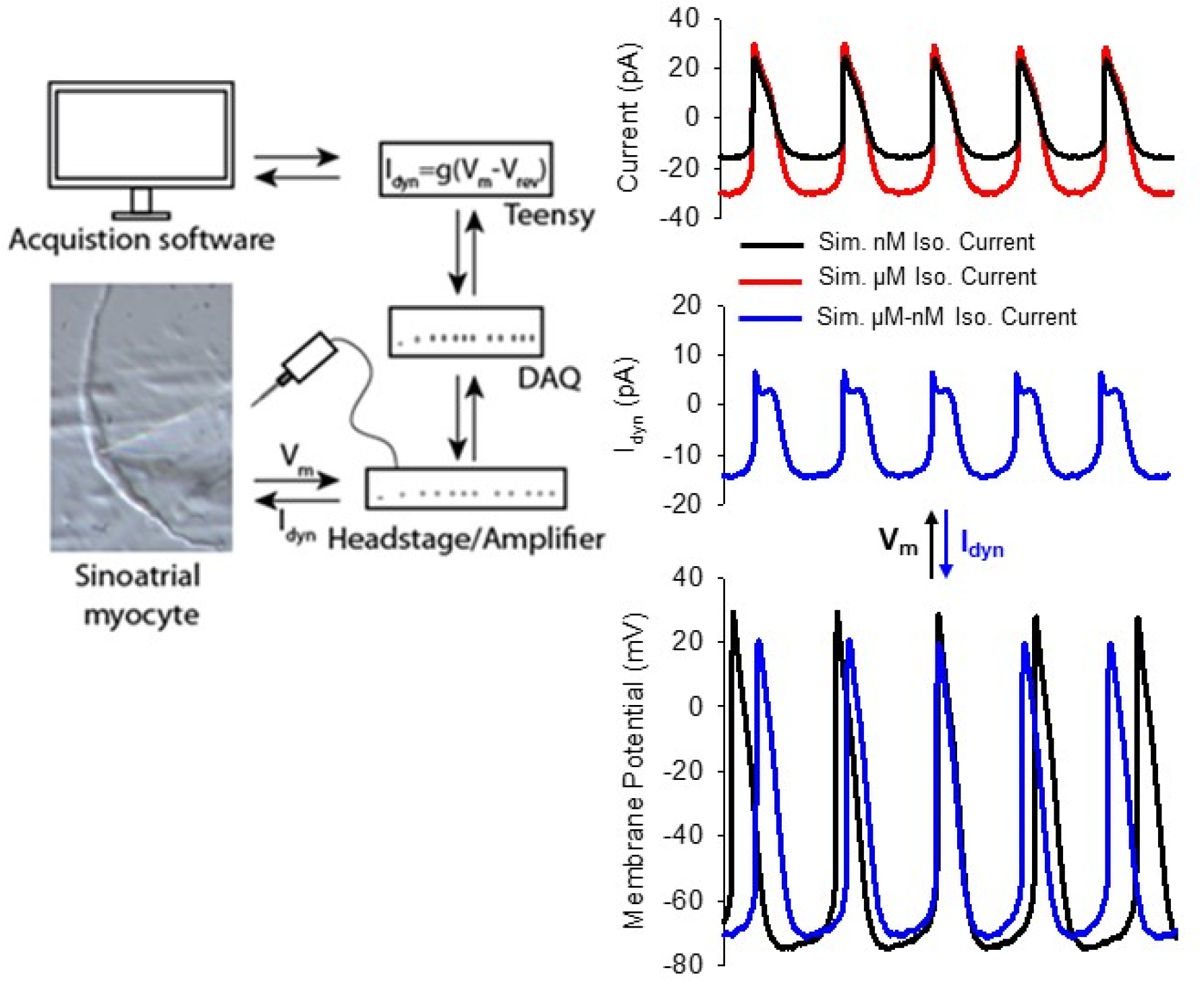
Dynamic Clamp Approach. **A:** Schematic of the dynamic clamp technique. Membrane voltages recorded from a patch-clamped SAM are used by a Teensy microprocessor to simulate currents using the 1 nM and 1 µM Iso models (*top right*). The calculated dynamic clamp current (I_dyn_), corresponding to the Iso-sensitive component of I_f_ (*middle right*), is then injected back into the SAM in a fast feedback loop and the resulting changes to AP firing are recorded (*bottom right*).

**Figure S3:**
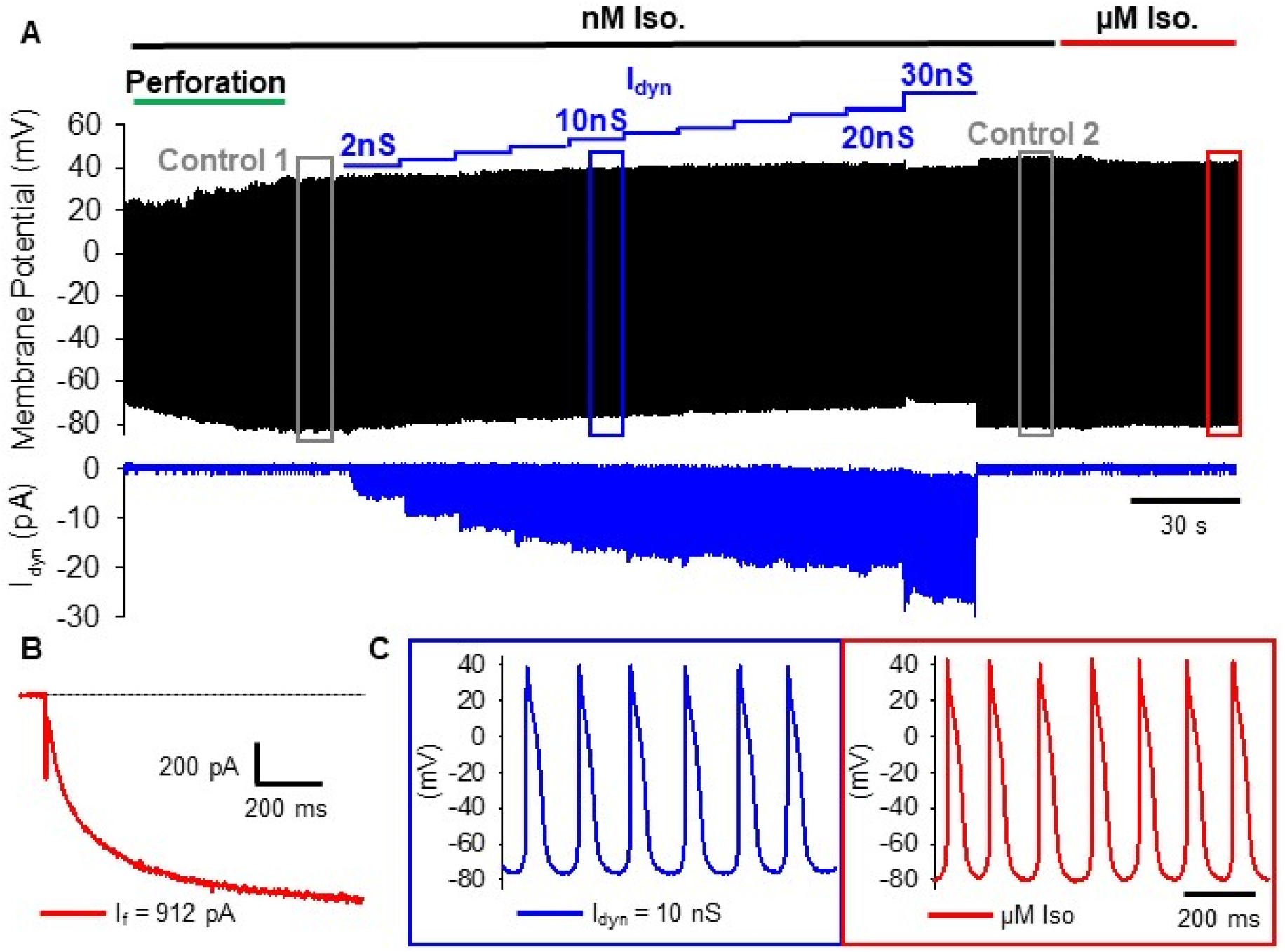
Cell-Scaled Dynamic Clamp Approach. **A-C:** Protocol for dynamic clamp recordings with simulated conductance scaled to endogenous I_f_. **A:** Spontaneous APs are recorded from a cell perfused with 1 nM Iso (*top*) and dynamic clamp current is monitored (*bottom*). After AP amplitude stabilizes following perforation (*green*), the set of control APs are recorded (*gray box*), after which the dynamic clamp circuit is switched on and the maximal I_f_ conductance used to calculate the injected current (I_dyn_) is increased in 2 nS steps (*blue*). Following the injected conductance series, the dynamic clamp circuit is switched off, a second set of control APs are recorded and the cell is perfused with 1 µM Iso to measure the full βAR-stimulated response (*red*). **B:** Following the control, dynamic current clamp, and Iso-stimulated recordings of APs in current-clamp mode, the amplifier is changed to voltage-clamp mode. 1 mM Ba^2+^ is perfused to block inward rectifier currents and the endogenous I_f_ conductance is calculated from the current elicited by a voltage-step to -130 mV where the current is >90% activated. In this example, conductance was calculated as 9.58 nS based on the 912 pA current and I_f_ reversal potential of -30 mV. **C:** The endogenous I_f_ is then used to post-hoc determine the dynamic clamp conductance for which the AP firing rate and waveform will be analyzed (*blue*).

## References

Alig, J., L. Marger, P. Mesirca, H. Ehmke, M.E. Mangoni, and D. Isbrandt. 2009. Control of heart rate by cAMP sensitivity of HCN channels. Proc. Natl. Acad. Sci. U. S. A.106:12189–12194. doi:10.1073/pnas.0810332106.

Baruscotti, M., A. Bucchi, C. Viscomi, G. Mandelli, G. Consalez, T. Gnecchi-Rusconi, N. Montano, K.R. Casali, S. Micheloni, A. Barbuti, and D. DiFrancesco. 2011. Deep bradycardia and heart block caused by inducible cardiac-specific knockout of the pacemaker channel gene Hcn4. Proc. Natl. Acad. Sci. U. S. A. 108:1705–1710. doi:10.1073/pnas.1010122108.

Breiman, L. 2001. Random Forests. Mach. Learn. 45:5–32. doi:10.1023/A:1010933404324.

Brown, H.F., D. Difrancesco, and S.J. Noble. 1979. How does adrenaline accelerate the heart? Nature. 280:235–236. doi:10.1038/280235a0.

Bucchi, A., A. Barbuti, D. Difrancesco, and M. Baruscotti. 2012. Funny Current and Cardiac Rhythm: Insights from HCN Knockout and Transgenic Mouse Models. Front. Physiol. 3:240. doi:10.3389/fphys.2012.00240.

Bucchi, A., M. Baruscotti, and D. DiFrancesco. 2002. Current-dependent Block of Rabbit Sino-Atrial Node If Channels by Ivabradine. J. Gen. Physiol. 120:1–13. doi:10.1085/jgp.20028593.

Bucchi, A., M. Baruscotti, R.B. Robinson, and D. DiFrancesco. 2007. Modulation of rate by autonomic agonists in SAN cells involves changes in diastolic depolarization and the pacemaker current. J. Mol. Cell. Cardiol. 43:39–48. doi:10.1016/j.yjmcc.2007.04.017.

Cai, Q., M. Lei, and H. Brown. 1995. Responses of guinea-pig SA node/atria to acetylcholine and adrenaline in the presence of blockers of If and IK,ACh. J. Physiol. 483:21P.

Desai, N. 2017. Microprocessor-based dynamic clamp.

DiFrancesco, D. 1982. Block and activation of the pace-maker channel in calf purkinje fibres: effects of potassium, caesium and rubidium. J. Physiol. 329:485–507.

DiFrancesco, D. 2010. The role of the funny current in pacemaker activity. Circ. Res. 106:434–446. doi:10.1161/CIRCRESAHA.109.208041.

Du, X.-J., X. Feng, X.-M. Gao, T.P. Tan, H. Kiriazis, and A.M. Dart. 2004. If channel inhibitor ivabradine lowers heart rate in mice with enhanced sympathoadrenergic activities. Br. J. Pharmacol. 142:107–112. doi:10.1038/sj.bjp.0705696.

Fenske, S., K. Hennis, R.D. Rötzer, V.F. Brox, E. Becirovic, A. Scharr, C. Gruner, T. Ziegler, V. Mehlfeld, J. Brennan, I.R. Efimov, A.G. Pauža, M. Moser, C.T. Wotjak, C. Kupatt, R. Gönner, R. Zhang, H. Zhang, X. Zong, M. Biel, and C. Wahl-Schott. 2020. cAMP-dependent regulation of HCN4 controls the tonic entrainment process in sinoatrial node pacemaker cells. Nat. Commun. 11:5555. doi:10.1038/s41467-020-19304-9.

Gao, Z., T.P. Rasmussen, Y. Li, W. Kutschke, O.M. Koval, Y. Wu, Y. Wu, D.D. Hall, M.A. Joiner, X.-Q. Wu, P.D. Swaminathan, A. Purohit, K. Zimmerman, R.M. Weiss, K.D. Philipson, L. Song, T.J. Hund, and M.E. Anderson. 2013. Genetic inhibition of Na+-Ca2+ exchanger current disables fight or flight sinoatrial node activity without affecting resting heart rate. Circ. Res. 112:309–317. doi:10.1161/CIRCRESAHA.111.300193.

Haechl, N., J. Ebner, K. Hilber, H. Todt, and X. Koenig. 2019. Pharmacological Profile of the Bradycardic Agent Ivabradine on Human Cardiac Ion Channels. Cell. Physiol. Biochem. Int. J. Exp. Cell. Physiol. Biochem. Pharmacol. 53:36–48. doi:10.33594/000000119.

Hagiwara, N., H. Irisawa, and M. Kameyama. 1988. Contribution of two types of calcium currents to the pacemaker potentials of rabbit sino-atrial node cells. J. Physiol. 395:233–253.

Harzheim, D., K.H. Pfeiffer, L. Fabritz, E. Kremmer, T. Buch, A. Waisman, P. Kirchhof, U.B. Kaupp, and R. Seifert. 2008. Cardiac pacemaker function of HCN4 channels in mice is confined to embryonic development and requires cyclic AMP. EMBO J. 27:692–703. doi:10.1038/emboj.2008.3.

Hennis, K., R.D. Rötzer, C. Piantoni, M. Biel, C. Wahl-Schott, and S. Fenske. 2021. Speeding Up the Heart? Traditional and New Perspectives on HCN4 Function. Front. Physiol. 12:669029. doi:10.3389/fphys.2021.669029.

Herrmann, S., J. Stieber, G. Stöckl, F. Hofmann, and A. Ludwig. 2007. HCN4 provides a “depolarization reserve” and is not required for heart rate acceleration in mice. EMBO J. 26:4423–4432. doi:10.1038/sj.emboj.7601868.

Hoesl, E., J. Stieber, S. Herrmann, S. Feil, E. Tybl, F. Hofmann, R. Feil, and A. Ludwig. 2008. Tamoxifen-inducible gene deletion in the cardiac conduction system. J. Mol. Cell. Cardiol. 45:62–69. doi:10.1016/j.yjmcc.2008.04.008.

Keith, A., and M. Flack. 1907. The Form and Nature of the Muscular Connections between the Primary Divisions of the Vertebrate Heart. J. Anat. Physiol. 41:172–189.

Lai, M.H., Y. Wu, Z. Gao, M.E. Anderson, J.E. Dalziel, and A.L. Meredith. 2014. BK channels regulate sinoatrial node firing rate and cardiac pacing in vivo. Am. J. Physiol. Heart Circ. Physiol. 307:H1327–1338. doi:10.1152/ajpheart.00354.2014.

Lakatta, E.G., and D. DiFrancesco. 2009. JMCC Point-Counterpoint. J. Mol. Cell. Cardiol. 47:157–170. doi:10.1016/j.yjmcc.2009.03.022.

Lakatta, E.G., V.A. Maltsev, and T.M. Vinogradova. 2010. A coupled SYSTEM of intracellular Ca2+ clocks and surface membrane voltage clocks controls the timekeeping mechanism of the heart’s pacemaker. Circ. Res. 106:659–673. doi:10.1161/CIRCRESAHA.109.206078.

Larson, E.D., J.R.S. Clair, W.A. Sumner, R.A. Bannister, and C. Proenza. 2013. Depressed pacemaker activity of sinoatrial node myocytes contributes to the age-dependent decline in maximum heart rate. Proc. Natl. Acad. Sci. 110:18011–18016.

Lees-Miller, J.P., J. Guo, J.R. Somers, D.E. Roach, R.S. Sheldon, D.E. Rancourt, and H.J. Duff. 2003. Selective Knockout of Mouse ERG1 B Potassium Channel Eliminates IKr in Adult Ventricular Myocytes and Elicits Episodes of Abrupt Sinus Bradycardia. Mol. Cell. Biol. 23:1856–1862. doi:10.1128/MCB.23.6.1856-1862.2003.

Lei, M., S.A. Jones, J. Liu, M.K. Lancaster, S.S.-M. Fung, H. Dobrzynski, P. Camelliti, S.K.G. Maier, D. Noble, and M.R. Boyett. 2004. Requirement of neuronal-and cardiac-type sodium channels for murine sinoatrial node pacemaking. J. Physiol. 559:835–848. doi:10.1113/jphysiol.2004.068643.

Leitch, S.P., H.F. Brown, and D.J. Paterson. 1993. Effect of caesium and β-adrenergic agonists on the rate of spontaneous activity in an isolated rabbit sino-atrial node preparation. J. Physiol. 459:86P.

Liao, Z., D. Lockhead, E.D. Larson, and C. Proenza. 2010. Phosphorylation and modulation of hyperpolarization-activated HCN4 channels by protein kinase A in the mouse sinoatrial node. J. Gen. Physiol. 136:247–258. doi:10.1085/jgp.201010488.

Lipsius, S.L., and D.M. Bers. 2003. Cardiac pacemaking: If vs. Ca2+, is it really that simple? J. Mol. Cell. Cardiol. 35:891–893. doi:10.1016/S0022-2828(03)00184-6.

Lucot, J.B., N. Jackson, I. Bernatova, and M. Morris. 2005. Measurement of plasma catecholamines in small samples from mice. J. Pharmacol. Toxicol. Methods. 52:274–277. doi:10.1016/j.vascn.2004.11.004.

Mesirca, P., J. Alig, A.G. Torrente, J.C. Müller, L. Marger, A. Rollin, C. Marquilly, A. Vincent, S. Dubel, I. Bidaud, A. Fernandez, A. Seniuk, B. Engeland, J. Singh, L. Miquerol, H. Ehmke, T. Eschenhagen, J. Nargeot, K. Wickman, D. Isbrandt, and M.E. Mangoni. 2014. Cardiac arrhythmia induced by genetic silencing of “funny” (f) channels is rescued by GIRK4 inactivation. Nat. Commun. 5:4664. doi:10.1038/ncomms5664.

Messan, F., A. Tito, P. Gouthon, K.B. Nouatin, I.B. Nigan, A.S. Blagbo, J. Lounana, and J. Medelli. 2017. Comparison of Catecholamine Values Before and After Exercise-Induced Bronchospasm in Professional Cyclists. Tanaffos. 16:136–143.

Milano, A., A.M.C. Vermeer, E.M. Lodder, J. Barc, A.O. Verkerk, A.V. Postma, I.A.C. van der Bilt, M.J.H. Baars, P.L. van Haelst, K. Caliskan, Y.M. Hoedemaekers, S. Le Scouarnec,

R. Redon, Y.M. Pinto, I. Christiaans, A.A. Wilde, and C.R. Bezzina. 2014. HCN4 mutations in multiple families with bradycardia and left ventricular noncompaction cardiomyopathy. J. Am. Coll. Cardiol. 64:745–756. doi:10.1016/j.jacc.2014.05.045.

Moosmang, S., M. Biel, F. Hofmann, and A. Ludwig. 1999. Differential distribution of four hyperpolarization-activated cation channels in mouse brain. Biol. Chem. 380:975–980. doi:10.1515/BC.1999.121.

Pedregosa, F., G. Varoquaux, A. Gramfort, V. Michel, B. Thirion, O. Grisel, M. Blondel, P. Prettenhofer, R. Weiss, V. Dubourg, J. Vanderplas, A. Passos, D. Cournapeau, M. Brucher, M. Perrot, and É. Duchesnay. 2011. Scikit-learn: Machine Learning in Python. J. Mach. Learn. Res. 12:2825–2830.

Peters, C.H., P.W. Liu, S. Morotti, S.C. Gantz, E. Grandi, B.P. Bean, and C. Proenza. 2021. Bidirectional flow of the funny current (If) during the pacemaking cycle in murine sinoatrial node myocytes. Proc. Natl. Acad. Sci. 118. doi:10.1073/pnas.2104668118.

Pichavaram, P., W. Yin, K.W. Evanson, J.H. Jaggar, and S. Mancarella. 2018. Elevated plasma catecholamines functionally compensate for the reduced myogenic tone in smooth muscle STIM1 knockout mice but with deleterious cardiac effects. Cardiovasc. Res. 114:668–678. doi:10.1093/cvr/cvy015.

Protze, S.I., J. Liu, U. Nussinovitch, L. Ohana, P.H. Backx, L. Gepstein, and G.M. Keller. 2017. Sinoatrial node cardiomyocytes derived from human pluripotent cells function as a biological pacemaker. Nat. Biotechnol. 35:56–68. doi:10.1038/nbt.3745.

Ravagli, E., A. Bucchi, C. Bartolucci, M. Paina, M. Baruscotti, D. DiFrancesco, and S. Severi. 2016. Cell-specific Dynamic Clamp analysis of the role of funny If current in cardiac pacemaking. Prog. Biophys. Mol. Biol. 120:50–66. doi:10.1016/j.pbiomolbio.2015.12.004.

Rickert, C., and C. Proenza. 2017. ParamAP: Standardized Parameterization of Sinoatrial Node Myocyte Action Potentials. Biophys. J. 113:765–769. doi:10.1016/j.bpj.2017.07.001.

Rickert, C., and C. Proenza. 2019a. pyClamp: A live graphical user interface for the dyClamp sketch. doi:10.5281/zenodo.2825278.

Rickert, C., and C. Proenza. 2019b. dyClamp: A real-time dynamic clamp sketch for the pyClamp interface. doi:10.5281/zenodo.2824830.

Rickert, C., and C. Proenza. 2021. christianrickert/ParamAP: Third release. doi:10.5281/zenodo.5250113.

Severi, S., M. Fantini, L.A. Charawi, and D. DiFrancesco. 2012. An updated computational model of rabbit sinoatrial action potential to investigate the mechanisms of heart rate modulation. J. Physiol. 590:4483–4499. doi:10.1113/jphysiol.2012.229435.

Sharpe, E.J., J.R.S. Clair, and C. Proenza. 2016. Methods for the Isolation, Culture, and Functional Characterization of Sinoatrial Node Myocytes from Adult Mice. JoVE J. Vis. Exp. e54555. doi:10.3791/54555.

Sharpe, E.J., E.D. Larson, and C. Proenza. 2017. Cyclic AMP reverses the effects of aging on pacemaker activity and If in sinoatrial node myocytes. J. Gen. Physiol. jgp.201611674. doi:10.1085/jgp.201611674.

Sosunov, E.A., and E.P. Anyukhovsky. 2012. Differential effects of ivabradine and ryanodine on pacemaker activity in canine sinus node and purkinje fibers. J. Cardiovasc. Electrophysiol. 23:650–655. doi:10.1111/j.1540-8167.2011.02285.x.

Stieber, J., S. Herrmann, S. Feil, J. Löster, R. Feil, M. Biel, F. Hofmann, and A. Ludwig. 2003. The hyperpolarization-activated channel HCN4 is required for the generation of pacemaker action potentials in the embryonic heart. Proc. Natl. Acad. Sci. U. S. A. 100:15235–15240. doi:10.1073/pnas.2434235100.

Stieber, J., K. Wieland, G. Stöckl, A. Ludwig, and F. Hofmann. 2006. Bradycardic and Proarrhythmic Properties of Sinus Node Inhibitors. Mol. Pharmacol. 69:1328–1337. doi:10.1124/mol.105.020701.

Swedberg, K., M. Komajda, M. Böhm, J.S. Borer, I. Ford, A. Dubost-Brama, G. Lerebours, and L. Tavazzi. 2010. Ivabradine and outcomes in chronic heart failure (SHIFT): a randomised placebo-controlled study. The Lancet. 376:875–885. doi:10.1016/S0140-6736(10)61198-1.

Torrente, A.G., R. Zhang, H. Wang, A. Zaini, B. Kim, X. Yue, K.D. Philipson, and J.I. Goldhaber. 2017. Contribution of small conductance K+ channels to sinoatrial node pacemaker activity: insights from atrial-specific Na+ /Ca2+ exchange knockout mice. J. Physiol. 595:3847–3865. doi:10.1113/JP274249.

Verkerk, A.O., G.S.C. Geuzebroek, M.W. Veldkamp, and R. Wilders. 2012. Effects of Acetylcholine and Noradrenalin on Action Potentials of Isolated Rabbit Sinoatrial and Atrial Myocytes. Front. Physiol. 3. doi:10.3389/fphys.2012.00174.

Zhang, H., and M. Vassalle. 2003. Mechanisms of adrenergic control of sino-atrial node discharge. J. Biomed. Sci. 10:179–192. doi:10.1007/BF02256053.

